# Discovery of a novel stereospecific β-hydroxyacyl-CoA lyase/thioesterase shared by three metabolic pathways in *Mycobacterium tuberculosis*

**DOI:** 10.1101/322404

**Authors:** Hua Wang, Alexander A. Fedorov, Elena V. Fedorov, Deborah M. Hunt, Angela Rodgers, Acely Garza-Garcia, Jeffrey B. Bonanno, Steven C. Almo, Luiz Pedro S. de Carvalho

## Abstract

The vast number of poorly characterised enzymes in *Mycobacterium tuberculosis* (Mtb) is one of the key barriers precluding a better understanding of the biology that underpins pathogenesis. Here, we investigated the Mtb orphan enzyme Rv2498c to delineate its physiological role. Our results from *in vitro* enzymatic assays, phylogenetic analysis, X-ray crystallography and *in vivo* Mtb experiments, de-orphan Rv2498c as a multi-functional β-hydroxyacyl-CoA lyase/thioesterase (β-HAClyase/thioesterase) that participates in three different metabolic pathways: L-leucine catabolism, itaconate dissimilation, and glyoxylate shunt. Moreover, the deletion of the *rv2498c* gene from the Mtb genome resulted in attenuation in the mouse model compared to infection with the parent strain. To the best of our knowledge, this is the first report of an (*R*)-3-hydroxyl-3-methylglutaryl-CoA for leucine catabolism and an itaconate-specific resistance mechanism in Mtb.

## Introduction

*Mycobacterium tuberculosis* (Mtb) is the aetiological agent of tuberculosis (TB). TB is one of the leading causes of death worldwide, accounting for approximately 1.3 million deaths in 2016, and is estimated to infect one-fourth of the global population.[1, 2] The pathogenicity, physiological resiliency and plasticity of Mtb are notably complex, with humans serving as their only known reservoir, highlighting the effect of niche adaptation on pathogen evolution.[3, 4] However, Mtb biology remains largely unexplored and over half of the enzymes in the proteome remain ‘orphan enzymes’, *i.e*. without a defined catalytic activity.[5, 6] To date, delineating the *in vitro* activities and *in vivo* functional roles of hundreds of poorly characterised enzymes has been a significant challenge.[7] This difficulty is in part due to the unconventional nutrient assimilation of Mtb and its ability to persist in different metabolic states.[8–12] Without an adequate understanding of the fundamental biology underpinning infection and the associated metabolic networks, we continue to generate drug candidates that fail in eradicating the pathogen. Thus, a deeper understanding of the Mtb biology is pivotal to tackle the TB pandemic.

Mtb protein Rv2498c is currently annotated as the β-subunit (CitE) of the trimeric prokaryotic citrate lyase complex involved in tricarboxylic acid (TCA) cycle cataplerosis.[13–15] However, the genes encoding the associated α- and γ- subunits of the citrate lyase complex are absent in the Mtb genome, pointing to a different but related function.[13, 16] *Bona fide* CitE enzymes catalyse the cleavage of (3*S*)-citryl-CoA to acetate and oxaloacetate (EC 4.1.3.34), thus Rv2498c is most probably also a short-chain acyl-CoA lyase but of unknown substrate specificity and physiological role. Rv2498c shares sequence and structural similarity with bacterial lyases that act on analogues of β-hydroxyl-positioned CoA-thioesters (*e.g*. 3-hydroxy-3-methylglutaryl-CoA (HMG-CoA), malyl-CoA, β-methylmalyl-CoA, and citramalyl-CoA).[17–21] These enzymes are involved in a variety of pathways including leucine catabolism, itaconate dissimilation, glyoxylate shunt, acetate assimilation, carbon dioxide fixation, C1 assimilation *via* the serine cycle and *via* the ethylmalonyl-CoA pathway, C2 assimilation *via* the ethylmalonyl-CoA pathway, and the methylaspartate pathway.

Here we report on the functional characterisation of the Mtb enzyme Rv2498c as a stereospecific multi-functional β-hydroxyacyl-CoA lyase/thioesterase (β-HAClyase/thioesterase) participating in three metabolic pathways: (1) L-leucine catabolism; (2) itaconate dissimilation; and (3) glyoxylate shunt. To the best of our knowledge, this is the first report that shows that Mtb can dissimilate itaconate, that Mtb leucine catabolism solely involves the (*R*)-HMG-CoA, and that Mtb has a second malate synthase. Additionally, we present Rv2498c ligand-bound crystal structures revealing the molecular foundations underlying the stereospecificity and of the β-hydroxyacyl-CoA thioester carbon-carbon cleavage. Overall, these findings offer new insights on the Mtb biology and physiological plasticity highlighting an example of a multi-functional enzyme.

## Results

### Rv2498c has (*R*)-HMG-CoA lyase activity *in vitro* and *ex vivo*

*Rv2498c* is reported to encode the β-subunit CitE-like of citrate lyase complex.[7, 13] Inconsistent with this annotation, the associated α- and γ-subunits needed to form the functional citrate lyase complex appear to be absent in Mtb. This disconnect prompted us to screen a diverse panel of commercially available CoA-thioesters using an UV-Vis HPLC-based assay (**Supplementary Table 1**).

In this screening, incubation of Rv2498c with HMG-CoA generated a product with a retention time similar to acetyl-CoA (Ac-CoA) and consumed exactly half of the HMG-CoA racemic mixture, suggesting absolute stereospecificity (**Fig 1a**); the stereoisomer (*S*)-HMG-CoA is a known metabolic intermediate in the leucine catabolic pathway, which is incompletely annotated in Mtb, while the role of the stereoisomer (*R*)-HMG-CoA is not well understood.[22] To unambiguously determine the stereospecificity of Rv2498c for HMG-CoA, we eliminated either the (*R*)- or (*S*)-isomer of HMG-CoA by taking advantage of the known stereospecific HMG-CoA lyases, *P. aeruginosa* PA0883 and PA2011.[18] Interestingly, we found that Rv2498c is a lyase specific for the (*R*)-HMG-CoA. Based on the observation that Ac-CoA is one product, the other likely product of C-C bond cleavage is acetoacetate (AAc), which is not visible in our UV-Vis HPLC assay. To directly detect the formation of AAc, we analysed the reaction products by ^1^H-NMR spectroscopy and observed the formation of Ac-CoA and AAc from half of the (*R*/*S*)-HMG-CoA in the reaction mixture (**Fig. 1b**). These results demonstrate the stereospecific (*R*)-HMG-CoA lyase activity of Rv2498c.

**Figure 1.**
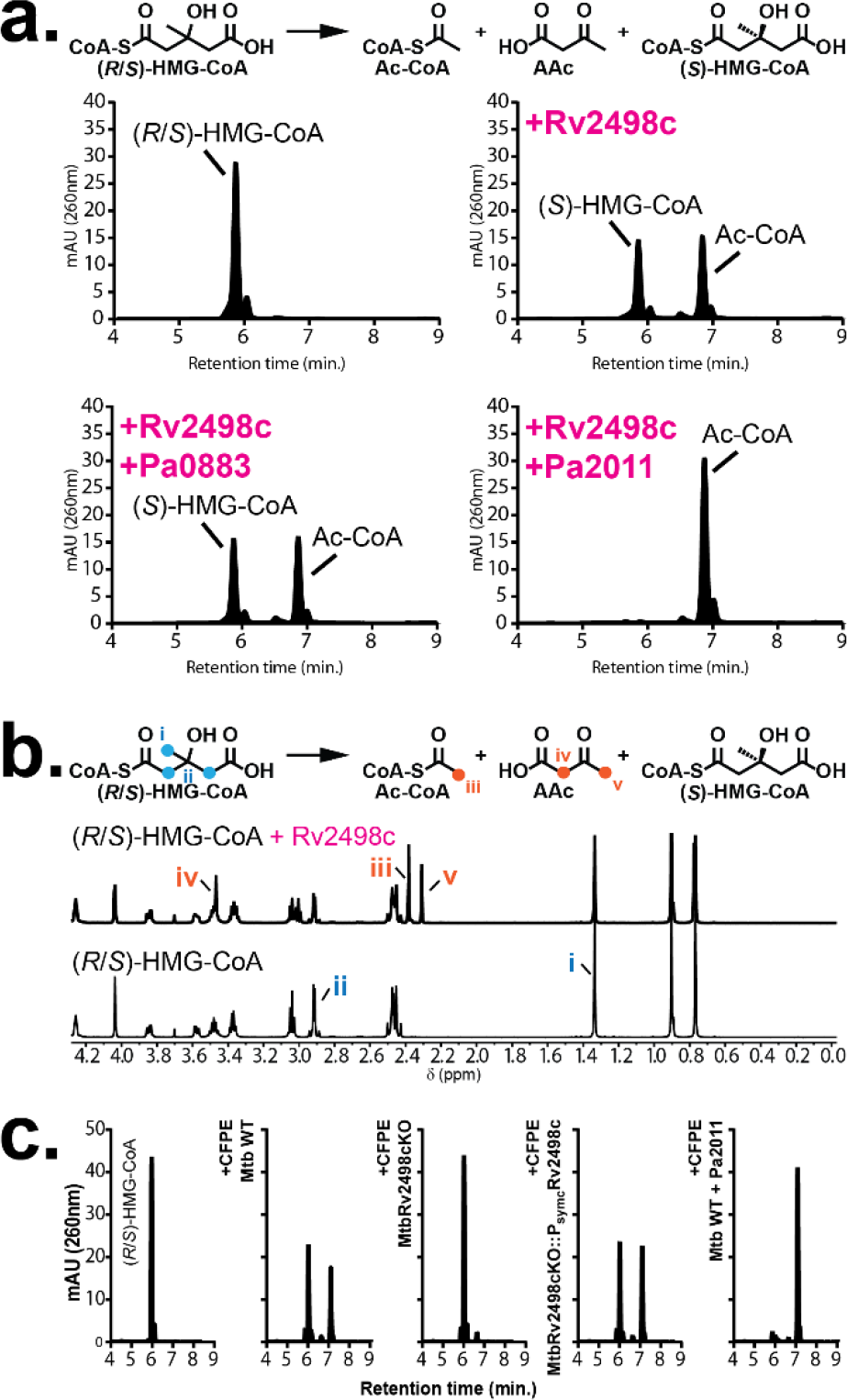
Rv2498c is a stereospecific lyase that cleaves (*R*)-HMG-CoA to produce acetyl-CoA and acetoacetate. **a**, HPLC chromatograms of 3-hydroxy-3-methylglutaryl-CoA racemic mixture (HMG-CoA) incubated with recombinant Rv2498c. The (*R*)-HMG-CoA lyase stereospecificity of Rv2498c is revealed when adding to the reaction either (*R*)-HMG-CoA-specific lyase Pa0883, or (*S*)-HMG-CoA-specific lyase Pa2011. Rv2498c with Pa0883 only consumed half of HMG-CoA, while Rv2498c with Pa2011 consumed all HMG-CoA. **b,** Comparison between the ^1^H NMR spectra of HMG-CoA incubated with or without recombinant Rv2498c. Peaks assigned to acetyl-CoA −CH_3_ (iii) group and acetoacetate −CH_3_ (v) and −CH_2_- (iv) groups are only observed in the spectrum with Rv2498c **c,** HPLC chromatograms of HMG-CoA incubated with Mtb CFPEs with or without (*S*)-HMG-CoA-specific lyase Pa2011. Degradation of (*R*)-HMG-CoA is not observed in the *rv2498c*KO CFPE chromatogram indicating that Rv2498c is needed for the activity.

To validate the results obtained using purified recombinant Rv2498c, we investigated the stereo-chemical course of HMG-CoA degradation in cell-free protein extracts (CFPE) from Mtb H37Rv (parent) and from an *rv2498c*-knockout strain (*rv2498c*KO). Consistent with the results using recombinant Rv2498c, parent CFPE, but not *rv2498c*KO CFPE, degraded (*R*)-HMG-CoA (**Fig. 1c**). These results confirm the (*R*)-specific stereo-chemical course of HMG-CoA degradation in Mtb and demonstrate that Rv2498c is both necessary and sufficient for breakdown of the (*R*)-HMG-CoA in CFPE.

### Rv2498c (*S*)-citramalyl-CoA bound structure reveals the molecular basis for stereospecificity

To investigate the molecular basis of Rv2498c substrate stereospecificity, we determined the X-ray crystal structure of Rv2498c in variety of liganded states (**Supplementary Table 2**). All structures showed the same trimeric arrangement of protomers previously described for the unliganded structure and for the oxaloacetate- and Mg^2+^-bound structure.[13] But in contrast to the previously reported structures, the C-terminus (50 residues) is well ordered, forming an α-helix/β-hairpin/α-helix motif that packs against the surface of the neighbouring protomer, capping its active site (**Fig. 2a**). The fact that the C-terminus is ordered in these ligand-bound structures is consistent with earlier observations that the position and organisation of the C-terminus might depend on the occupancy of the active site (**Fig. 2b**).[20]

**Figure 2.**
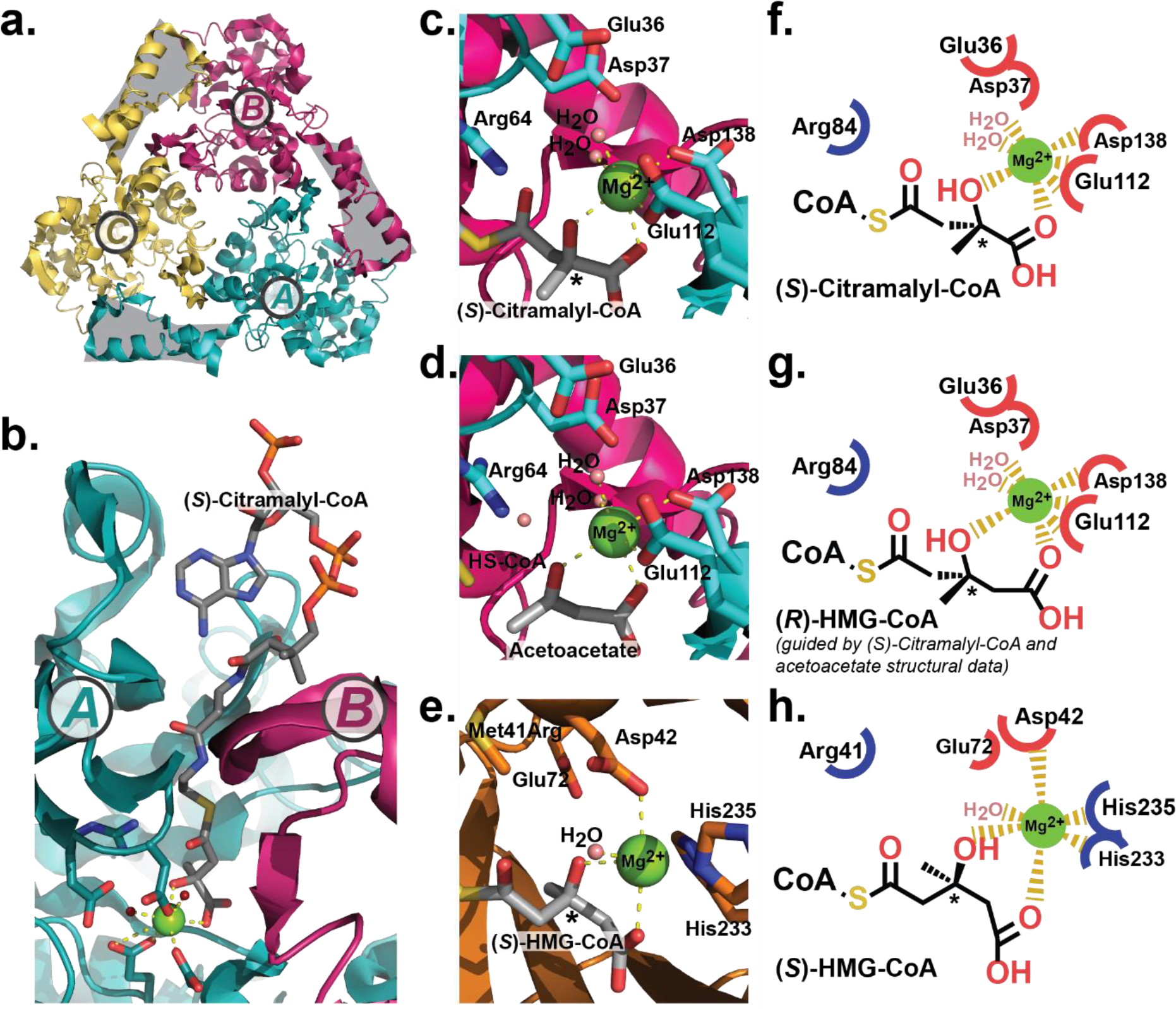
Ligand-bound X-ray 3D structures of Rv2498c reveal that metal coordination is the basis of stereoselectivity. **a,** Ribbon representation of trimeric Rv2498c with protomers A, B, and C (PDB: 6CJ4). The structured C-terminal domain is highlighted in grey. **b,** Ribbon and stick representation of Rv2498c bound to (*S*)-citramalyl-CoA (PDB: 6AQ4), highlighting the contribution of the C-terminal domain of protomer B (magenta) to the binding pocket of protomer A (cyan). **c,** Close-up view of the catalytic site of (*S*)-citramalyl-CoA-bound Rv2498c. The β-hydroxyl group chelation of the Mg^2+^ ion favours the *S*-conformation of citramalyl-CoA. **d,** Close-up view of the catalytic site of acetoacetate-bound Rv2498c (PDB: 6AS5). Acetoacetate, the HMG-CoA elimination product retains the keto-acid conformation. A similar view of the catalytic site of human (*S*)-HMG-CoA lyaseR41M mutant (PDB: 3MP5) bound to (*S*)-HMG-CoA is shown for comparison (**e**).**f**-**h,** Ligplot-type schematic diagrams of the catalytic sites of the experimental structure of Rv2498c bound to (*S*)-citramalyl-CoA (**f**), a model of Rv2498c bound to (*R*)-HMG-CoA (**g**), and a model of the human (*S*)-HMG-CoA lyase WT bound to (*S*)-HMG-CoA (**h**). The stereoselectivity between (*R*)- and (*S*)-HMG-CoA conformations depends on the geometry of the Mg^2+^ chelation by the β-hydroxyl group of the CoA ligand as shown in (**c**) and (**e**). Representation of (*R*)-HMG-CoA in Rv2498c (**g**) was modelled based on the reaction substrate (*S*)-citramalyl-CoA (**c**) and the reaction product acetoacetate (**d**). Ribbon and stick representations generated with PyMOL. Atoms are coloured according to the CPK colouring scheme. ‘*’ denotes a chiral carbon centre.

Rv2498c has no sequence and modest structural similarity to the characterised family of TIM barrel (*S*)-HMG-CoA lyases that includes human HMG-CoA lyase (PDB: 3MP5) and the bacterial lyases from *Brucella melitensis*, *Bacillus subtillis* and *Pseudomonas aeruginosa* (PDB: 1YDN, 1YDO & 2FTP).[23] Instead, the three most closely related ligand-bound structures to Rv2498c are the human citramalyl-CoA lyase CLYBL (PDB: 5VXO) and the bacterial L-malyl-CoA/ β-methylmalyl-CoA lyases from *Rhodobacter sphaeroides* (PDB: 4L9Y) and from *Chloroflexus aurantiacus* (PDB: 4L80) with RMSDs of 2.35 (over 261 Cα atoms), 2.43 (over 266 Cα atoms) and 2.65 (over 266 Cα atoms), respectively.[20, 24–26] These three proteins were crystallised with propionyl-CoA in their active sites.

In the structures presented here of Rv2498c bound to (*S*)-citramalyl-CoA (PDB: 6AQ4) and to acetoacetate (PDB: 6AS5), the CoA moiety binds in a deep cleft at the base of which resides the active site Mg^2+^ ion. The magnesium ion is coordinated by the 3-hydroxyl group and by an unidentate interaction with the terminal carboxylate of the ligand, as well as by Glu112, Asp138 and two water molecules (**Fig. 2c, d**). The carboxylate of the substrate is further positioned by polar interactions with the backbone NH atoms of Ala136, Glu137 and Asp138, while the 3-hydroxyl group lies close to Arg64. The 3-methyl group of the ligand contacts a hydrophobic surface formed by Gly135 and the side-chain atoms of Met133 and Val181 (**Supplementary Fig. S1**). Of note, the structure with (*S*)-citramalyl-CoA bound was obtained from crystals that were soaked with pyruvate and Ac-CoA, therefore indicating that the conformation of Rv2498c in the crystals is catalytic active.

Because the citramalyl group observed in the active site was in the *S* configuration, which corresponds to the *R* configuration at the 3-position of HMG-CoA (**Supplementary Fig. S2**), we could use this structure to directly model the mode of binding of (*R*)-HMG-CoA. In this case, the additional methanediyl unit (C4) found in HMG-CoA is readily accommodated in the pocket and can support a less distorted octahedral coordination of the Mg^2+^ ion (the bite angle of the citramalyl group is ~70°; the modelled HMG moiety bite angle is ~85°). Likewise, Molprobity calculations revealed shape and charge complementarity between the protein and modelled substrate, consistent with the active site supporting energetically favourable interactions with (*R*)-HMG-CoA.[27] Attempts to model (*S*)-HMG-CoA into the active site generated electrostatically unfavourable interactions. Interestingly, with respect to the human (*S*)-HMG-CoA lyase (**Fig. 2e**, PDB: 3MP5), the differential positioning of the magnesium ion coordination with conserved Asp and Glu residues in the active site contributes to the stereoselectivity between the HMG-CoA stereoisomers (**Fig. 2f-h**).

### Phylogenetic analysis suggests Rv2498c might have alternative substrates

Aiming to conduct a comprehensive comparison of Rv2498c in the context of related enzymes, we constructed an amino acid sequence phylogenetic tree of the PFAM HpcH/HpaI aldolase/citrate lyase family (PF03328; **Fig. 3**).[28] The relationship within the family members is complex; the sequences group into more than twenty, presumably paralogous, clusters. Most of the clusters contain only bacterial sequences, but important Archaeal- and fungal-only groups are also present. The irregular taxonomic distribution of the bacterial branches suggests extensive lateral transfer. There is only one cluster that includes metazoan sequences, that of the itaconate-detoxifying CLYBL.[25] The Rv2498c cluster is actinobacterial-specific and contains only uncharacterised sequences (seventy-two in UnitProt). A variety of enzymatic activities has been described for PF03328 members, but the majority fall into two classes as it is evident from the tree: aldehyde-lyases (EC:4.1.2) and oxo-acid-lyases (EC:4.1.3); Rv2498c belongs to the latter.

**Figure 3.**
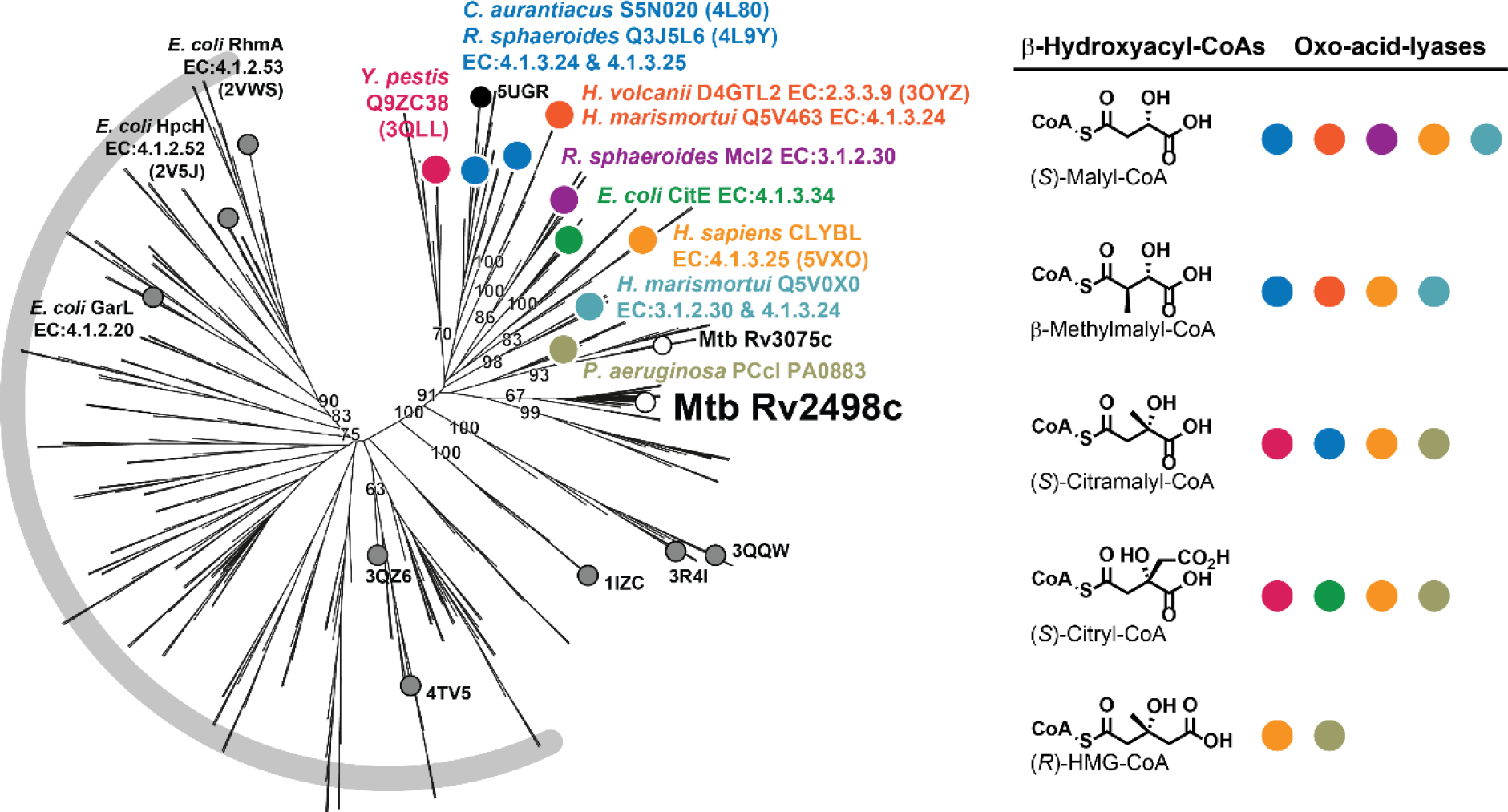
Phylogenetic tree of Rv2498c and related lyases/thioesterases pinpoints widespread substrate multiplicity. Neighbour-joining protein phylogenetic tree of the PFAM HpcH/Hpal aldolase/citrate lyase family (PF03328). The tree is mainly divided in two lobes: the aldehyde-lyases (EC: 4.1.2) shaded in grey and the oxo-acid-lyases (EC: 4.1.3). Mtb Rv2498c belongs to the latter, in a cluster of unknown substrate and function. The only two enzymes in the family in the Mtb genome are highlighted with white circles. Coloured circles denote experimentally characterised oxo-acid lyases shown with their respective β-hydroxyacyl-CoA substrates; PDB, UnitProtK and/or E.C. codes are provided. Characterised enzymes include the *Y. pestis* (*S*)-citramalyl-CoA lyase involved in itaconate dissimilation; *R. sphaeroides* (Mcl1) and *C. aurantiacus* (Mcl) L-malyl-CoA/β-methylmalyl-CoA lyases involved in CO_2_ fixation and acetate assimilation; archaeal *Haloferax volcanii* AceB and *Haloarcula marismortui* CitE1 L-malyl-CoA/β-methylmalyl-CoA lyases, involved in the methylaspartate cycle; *R. sphaeroides* (Mcl2) L-malyl-CoA thioesterase; *E. coli* CitE (*e.g*. P0A9I1) involved in TCA anaplerosis; human CLYBL (*S*)-citramalyl-CoA lyase involve in itaconate dissimilation; archaeal *H. marismortui* AceB L-malyl-CoA/β-methylmalyl-CoA lyase/thioesterase; and *P. aeruginosa* (*S*)-citramalyl-CoA lyase involved in itaconate dissimilation.

The similarity observed at the structural level between Rv2498c, CLYBL (*e.g*. UniProtKB: Q8N0X4), and the L-malyl-CoA/β-methylmalyl-CoA lyases from *R. sphaeroides* (Mcl1; Q3J5L6) and *C. aurantiacus* (Mcl; S5N020) is also evident at the level of the amino acid sequence. Other closely related groups are the *bona fide* CitE (*e.g*. P0A9I1), the archaeal L-malyl-CoA/β-methylmalyl-CoA lyases involved in the methylaspartate cycle (*e.g*. Q5V463), and the (*S*)-citramalyl-CoA lyases from *Yersinia pestis* (Q9ZC38) and *P. aeruginosa* (Pa0883; Q9I562).[18, 21] Coincidentally, another uncharacterised Mtb protein, Rv3075c, might be the orthologue of Pa0883.

Substrate and enzymatic function multiplicity are common in the PF03328 family. For example, *C. aurantiacus* Mcl catalyses (i) cleavage of (*S*)-malyl-CoA to acetyl-CoA and glyoxylate, (ii) condensation of glyoxylate of propionyl-CoA to form β-methylmalyl-CoA and (iii) cleavage of (*S*)-citramalyl-CoA into acetyl-CoA and pyruvate; while *Haloarcula marismortui* AceB (Q5V0X0) has been shown to condense acetyl-CoA with glyoxylate to form (*S*)-malyl-CoA and then to hydrolyse the thioester bond to form malate and free CoA.[19, 21, 29]

### Rv2498c also displays thioester hydrolase and malate synthase activities

Motivated by the results of the phylogenetic analysis (**Fig. 3**), we tested if (*S*)-citramalyl-CoA, (*S*)-malyl-CoA and/or β-methylmalyl-CoA are Rv2498c substrates. Because these molecules are not commercially available, these experiments were performed by analysing the reverse enzymatic reaction, *i.e*. synthase activity; driving Rv2498c to catalyse the formation of the CoA-thioesters from Ac-CoA or propionyl-CoA (Pro-CoA), and glyoxylate or pyruvate. (*S*)-Citramalyl-CoA, (*S*)-malyl-CoA, and β-methylmalyl-CoA, were all found to be substrates/ products. The identity of the products was confirmed by comparing retention times and liquid chromatography-mass spectrometry (LC-MS) spectra to those of the thioesters synthesised using *C. aurantiacus* Mcl [20, 30] (**Supplementary Fig. S3**, see **Online Methods**). The possibility of spontaneous or intramolecular general-base catalysed hydrolysis was ruled out for (*S*)-citramalyl-CoA and for (*S*)-malyl-CoA; but the results for β-methylmalyl-CoA were less clear, as formation of free CoA was observed overtime in the absence of Rv2498c.

Unexpectedly, Rv2498c was found to subsequently catalyse the hydrolysis of the thioester bond of (*S*)-malyl-CoA and β-methylmalyl-CoA both *in vitro* and in the parent vs *rv2498c*KO CFPE experimental set up (**Fig. 4c**, **Supplementary Fig. S4**), and is therefore able to synthesise malate and methylmalate by acting as a bifunctional (*S*)-malyl-CoA lyase/thioesterase as has been described for *H. marismortui* AceB.

**Figure 4.**
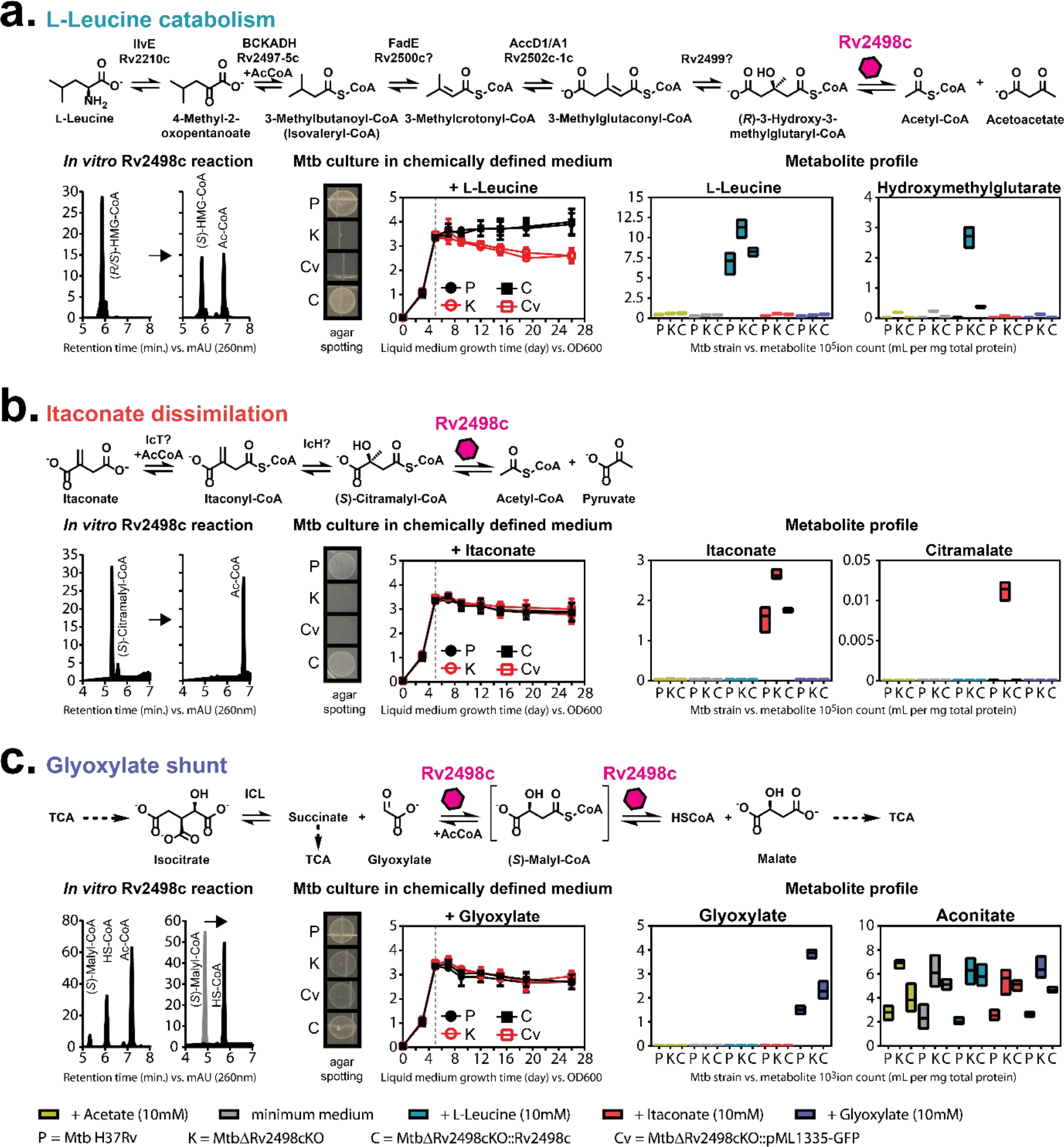
Rv2498c participates in L-leucine catabolism, itaconate dissimilation, and glyoxylate shunt in Mtb. **a-c** show reaction schemes of the proposed pathways accompanied with experimental data. ‘*In vitro* Rv2498c reaction’ shows HPLC chromatograms of Rv2498c reaction. ‘Mtb cultures in chemically defined medium’ shows solid and liquid growth conditions. The spotting cultures are growth with substrate (10 mM) after 31 days. The results are representative of three independent experiments. Liquid Mtb cultures are first grown in 7H9complete medium, then changed to chemically defined media with added substrate (10 mM) on day 5, indicated by the dotted line. The data are shown as mean values ± SD from three independent experiments. ‘Metabolite profile’ shows quantitative measurements of metabolite from filter culture growth on chemically defined media with added substrate (10 mM). The data are shown as mean values ± SD from three biological replicates. **a,** Rv2498c participation in Mtb L-leucine catabolism is demonstrated by (i) *in vitro* carbon-carbon bond cleavage of (*R*)-HMG-CoA to produce acetoacetate and acetyl-CoA (Ac-CoA), (ii) lack of growth of Mtb *rv2498c*KOs in L-leucine, and (iii) accumulation of leucine and HMG (from HMG-CoA hydrolysis) in Mtb *rv2498c*KO metabolite extract. **b,** Rv2498c participation in Mtb itaconate dissimilation is demonstrated by (i) *in vitro* carbon-carbon bond cleavage of (*S*)-citramalyl-CoA to produce pyruvate and Ac-CoA, (ii) absence of growth of Mtb *rv2498c*KOs in itaconate, and (iii) accumulation of itaconate and citramalate (from hydrolysed citramalyl-CoA) in Mtb *rv2498c*KO metabolite extract. **c,** Rv2498c participation in Mtb glyoxylate shunt is demonstrated by (i) *in vitro* malate synthase reaction from glyoxylate and Ac-CoA and hydrolysis of *in situ* (*S*)-malyl-CoA to produce malate and CoA, (ii) *in vitro* hydrolysis of synthesised (*S*)-malyl-CoA, (iii) reduced cell density of Mtb *rv2498c*KOs in glyoxylate, and (iv) accumulation of glyoxylate and aconitate in Mtb *rv2498c*KO metabolite extract. See **Supplementary Fig. S7**.

### Rv2498c is not an (*S*)-citryl-CoA lyase

The (*S*)-citryl-CoA lyase activity attributed to Rv2498c was directly tested in three different ways. (1) We monitored the reaction by UV-Vis HPLC using oxaloacetate and Ac-CoA as substrates. Production of citryl-CoA was not detected, instead the product was identified as citramalyl-CoA by retention time comparison with a standard and LC-MS (**Supplementary Fig. S2**). Formation of citramalyl-CoA was attributed to the short half-life of oxaloacetate, which spontaneously decarboxylated to form pyruvate.[31] (2) We monitored the reverse reaction by UV-Vis HPLC and by LC-MS using citrate and free CoA. No product formation was observed (**Supplementary Fig. S5**). (3) We synthesised (*S*)-citryl-CoA from inactivated citrate lyase, and confirmed that Rv2498c is unable to process (*S*)-citryl-CoA even in the presence of ATP (**Supplementary Fig. S5**).[32] In strict agreement with our results using recombinant enzyme, we found that (*S*)-citryl-CoA is readily hydrolysed by both parent and *rv2498c*KO CFPEs, confirming that Rv2498c is not the enzyme responsible for (*S*)-citryl-CoA hydrolysis (**Supplementary Fig. S5**).

In summary, we have shown that Rv2498c has β-hydroxyl-acyl-CoA lyase and thioesterase activities, and that (*R*)-HMG-CoA, (*S*)-citramalyl-CoA, (*S*)-malyl-CoA, and β-methylmalyl-CoA, but not (*S*)-citryl-CoA, are Rv2498c substrates. These results unambiguously establish that Rv2498c is not a CitE.

### Rv2498c participates in L-leucine and itaconate catabolism

We interrogated the role of Rv2498c in Mtb metabolism by comparing growth and targeted LC-MS metabolic profiles of *rv2498c*KO, parent and a *rv2498c*-complemented strain where the *rv2498c* gene is present elsewhere in the chromosome (*rv2498c*KO::*rv2498c*). These four strains were cultured in chemically defined media of composition similar to Middlebrook 7H10 for solid medium or 7H9 for liquid medium but with a single carbon source present. In accordance to our substrate specificity results using purified recombinant Rv2498c and CFPEs, we hypothesised that Rv2498c could be involved in at least three metabolic pathways: (1) L-leucine catabolism, in which (*R*)-HMG-CoA is a catabolic intermediate; (2) itaconate dissimilation, a process that involves (*S*)-citramalyl-CoA formation and; (3) glyoxylate shunt, where Rv2498c could function alongside or substitute for the *bona fide* malate synthase GlcB (Rv1837c).

Consistent with a role of Rv2498c in L-leucine metabolism, no growth was observed on solid or in liquid medium for the *rv2498c*KO strain in the presence of L-leucine as the sole carbon source. We inferred that this growth deficiency was likely the outcome of the absence of HMG-CoA lyase activity, resulting in the accumulation of HMG, limited production of Ac-CoA, and/or altered branched-lipid metabolism. Accordingly, in the presence of L-leucine as sole carbon source, *rv2498c*KO, but not the parent or the complemented strains, accumulated HMG (**Fig. 4a**), as well as other leucine catabolism intermediates such as methylcrotonoate, methylglutaconoate and hydroxymethylglutarate (**Supplementary Fig. S6**). We also observed a slight pH difference in the liquid medium of the Mtb *rv2498c*KO culture (pH 6.7) compared to the parent strain (pH 6.9) and attributed this acidification to an increased concentration HMG in the medium, as we could verify by LC-MS (**Supplementary Fig. S6**).

The phenotype of the cultures grown in medium with itaconate as sole carbon source was less clear-cut. A growth defect was observed for *rv2498c*KO compared to parent and complemented strains on solid medium, but none of the strains grew in liquid medium. Itaconate is a strong inhibitor of isocitrate lyase, the first enzyme of the glyoxylate shunt, and it is therefore considered to be an antimicrobial compound. The observed dissimilar growth pattern could be due to increased exposure to itaconate in liquid compared to solid medium (*e.g*. biofilm in solid medium reduces itaconate exposure). A pathway for itaconate degradation has not been described for Mtb, but in mammals and in some bacteria it is understood to proceed *via* activation of itaconate to itaconyl-CoA, stereospecific hydration to form (*S*)-citramalyl-CoA, and C-C bond cleavage to form pyruvate and acetyl-CoA.[18, 33] Consistent with the existence of this pathway in Mtb and with the involvement of Rv2498c in itaconate degradation, we observed accumulation of citramalate, the hydrolysis product of citramalyl-CoA, in the *rv2498c*KO strain when grown in itaconate as sole carbon source (**Fig. 4b**).

Similar to the cultures grown in itaconate liquid medium, no growth was observed for any of the strains in liquid medium containing glyoxylate, suggesting that Mtb is unable to use glyoxylate as a carbon source. However, *rv2498c*KO did show a decreased growth phenotype compared to the parent and the complemented strains on solid medium, perhaps because the cells are able to metabolise impurities in the agar. Nevertheless, when cells were grown with acetate or with glyoxylate as sole carbon source, we detected a discreet but reproducible accumulation of the TCA intermediate aconitate, a metabolite preceding the bifurcation of TCA and glyoxylate shunt, in *rv2498c*KO compared to parent and complemented strains. It is possible that in this instance, the *rv2498c*KO phenotype is obscured by the activity of the canonical malate synthase (**Fig. 4c**).

These *in vivo* experiments corroborate our *in vitro* and *ex vivo* biochemical data and offer further evidence that Rv2498c is a multi-functional enzyme participating in at least three metabolic pathways in Mtb: leucine catabolism, itaconate dissimilation, and glyoxylate shunt.

### *rv2498c*KO strain is attenuated in a mouse infection model

To test the impact of *rv2498c* deletion during infection, we carried out a low dose aerosol infection of C57BL/6J mice. The *rv2498c*KO strain showed a one log10 reduction in colony forming units (CFUs) in lung at day 28, 84, 112 post infection, compared to the parent strain (**Supplementary Fig. S8**). The decreased infection capacity observed for the *rv2498c*KO strain suggests that one or more of the activities associated with Rv2498c have a significant negative impact in the fitness of Mtb during experimental infection.

## Discussion

Our understanding of even the most well conserved and central metabolic pathways in Mtb is hampered by the prevalence in the genome of experimentally uncharacterised enzymes and enzymatic function database misannotations. Mtb Rv2498c is a striking example of an enzyme that eluded characterisation for over a decade, highlighting the intricacies and difficulties of enzyme functional assignment.

We found Rv2498c in a class of oxo-acid lyases with paradoxical features: characterised by promiscuity for substrates for carbon-carbon bond cleavage but absolute discrimination on the substrate stereochemistry. A systematic evaluation of substrate specificity revealed (*R*)-HMG-CoA, (*S*)-citramalyl-CoA, (*S*)-malyl-CoA, β-methylmalyl-CoA as substrates for Rv2498c.

We showed that Mtb catabolises L-leucine *via* an unprecedented use of (*R*)-HMG-CoA rather than the commonly attributed (*S*)-HMG-CoA isomer. Mtb lacking Rv2498c appears to suffer the same fate as humans with HMGCL-deficiency, the accumulation of HMG and keto acids in the leucine catabolic pathway.[34] In addition to L-leucine, we tested L-valine and L-isoleucine (the two other branched-chain amino acids) and observed no growth phenotypes between the parent and the *rv2498c*KO strains. These results agree with the known differences in catabolism of these three amino acids; HMG-CoA is only present in L-leucine degradation and not a shared metabolic intermediate in the catabolism of branched-chain amino acids.

We showed for the first time that Mtb can dissimilate itaconate, a hallmark macrophage metabolite synthesised after lipopolysaccharide activation during host inflammatory response to fight infection and also linked to Vitamin B12 deficiency.[25, 35, 36]. Itaconate is present in some prokaryotes, but its exact physiological function remains unclear.[18, 36–41] In contrast to the established toxic character of this molecule, our results suggest that Mtb can dissimilate itaconate, revealing a new facet where Mtb may be able to produce (*S*)-citramalyl-CoA from itaconate and to use Rv2498c to produce Ac-CoA and pyruvate from (*S*)-citramalyl-CoA, effectively using a host-derived antibacterial molecule as a nutrient source.[18, 33, 42]

We found that Rv2498c functions as a malate synthase by catalysing an aldol condensation followed by thioester hydrolysis and resulting in the formation of malate from glyoxylate and Ac-CoA, *via* a non-covalent malyl-CoA reaction intermediate. The physiological role for Rv2498c in the glyoxylate shunt is unclear as Mtb possesses a canonical malate synthase (GlcB, Rv1837c).[43] However, the first step in the shunt is also catalysed by two apparently redundant isocitrate lyases (Rv0467 and Rv1915-Rv1916). Such redundancy might be metabolically advantageous if the enzymes display different kinetic or regulatory properties. It is also possible that multi-functional enzymes, such as Rv2498c, are able to circumvent the presence of host-derived enzyme inhibitors of the canonical enzymes.[44, 45]

Our crystal structures for Rv2498c shed light into the molecular bases of substrate specificity and stereo-selectivity and revealed how the C-terminal domain interacts directly with the neighbouring CoA substrate, therefore playing an important role in substrate binding selectivity. We suggest that Rv2498c carbon-carbon cleavage of (*R*)-HMG-CoA follows a mechanism similar to that proposed for (*S*)-HMG-CoA lyase described by Fu *et al*. (**Supplementary Fig. S9**) and that the difference in the (*S*)- and (*R*)-stereospecificity for HMG-CoA can be partially attributed to the β-positioned hydroxyl group of the CoA-thioester coordination with the divalent metal.[46] We observed that the Rv2498c catalytic site has ordered water molecules, suggesting a catalytic mechanism for the carbon-carbon cleavage with water participation by shuttling protons and/or acting as the nucleophile in hydrolysis.[20, 46, 47] The lyase reaction likely proceeds *via* the reversal of standard retro-aldol condensation, as previously suggested.[46, 48]

In conclusion, we have defined the enzymatic activities of Rv2498c as a β-HAClyase/thioesterase and present full-length crystal structures of the protein, which revealed the details of its stereospecificity for (*R*)-HMG-CoA and the involvement of the C-terminal domain in substrate binding. Our results demonstrate for the first time that Mtb possesses (i) an (*R*)-specific HMG-CoA L-leucine catabolic pathway, (ii) the capability to dissimilate itaconate, and (iii) a secondary malate synthase. Importantly, deletion of *rv2498c* from the Mtb genome led to a defect during murine infection, indicating that one or more of its functions are involved in survival inside the host.

## Author contributions

In vitro experiments: H.W. Mtb experiments: H.W., D.M.H. Phylogenetic analysis: A.G.G. X-ray crystallography: A.A.F., E.V.F., J.B.B., S.C.A. LC-MS experiments: H.W. Mouse experiments: A.R. Data analysis: H.W., A.G.G., A.A.F., E.V.F., A.R., J.B.B., S.C.A., L.P.S.C. Manuscript preparation: H.W., A.G.G., J.B.B., S.C.A., L.P.S.C.

## ACKNOWLEDGMENTS

We thank Dr Geoff Kelly (MRC Biomedical NMR Centre) for assistance with NMR spectroscopy. NMR data were recorded in the MRC Biomedical NMR Centre at the Francis Crick Institute, which receives core funding from Cancer Research UK (FC001029); the Medical Research Council (FC001029); and the Wellcome Trust (FC001029). Work in LPSC’s laboratory was supported by the Francis Crick Institute, which receives its core funding from Cancer Research UK (FC001060), the UK Medical Research Council (FC001060), the Wellcome Trust (FC001060). LPSC’s laboratory also acknowledges funds from a Wellcome Trust New Investigator Award (104785/B/14/Z). This research used resources of the Advanced Photon Source, a U.S. Department of Energy (DOE) Office of Science User Facility operated for the DOE Office of Science by Argonne National Laboratory under Contract No. DE-AC02-06CH11357. Use of the Lilly Research Laboratories Collaborative Access Team (LRL-CAT) beamline at Sector 31 of the Advanced Photon Source was provided by Eli Lilly Company, which operates the facility. The Einstein Crystallographic Core X-Ray diffraction facility is supported by NIH Shared Instrumentation Grant S10 OD020068, which we gratefully acknowledge. SCA acknowledges support from the US National Institutes of Health (P01 GM118303).

## ONLINE METHODS

### Materials

All biological and chemical reagents were purchased from Sigma Aldrich or Fisher Scientific, unless stated otherwise. The following reagent was obtained through BEI Resources, NIAID, NIH: *Mycobacterium tuberculosis*, strain H37Rv, purified phthiocerol dimycocerosate (PDIM), NR-20328. Minimal medium was prepared in-house. *pMC2m* was a gift from Sabine Ehrt (Addgene Plasmid #17970)[1].

### Cloning of *rv2498c* gene

*rv2498c* gene was amplified from Mtb H37Rv genomic DNA by PCR (5’-GCCATATGGCACACCACCACCACCACCACATGAACCTGCGTGCCGCCGG-3’, 5’-GCAAGCTTTCATTCGGAGGTGGCTTCCC-3’), and ligated into *NdeI* and *HindIII* sites of pET-23a(+).

### Heterologous expression of Rv2498c, PA0883, PA0883, and MMClyase

*E.coli* BL21(DE3) cells were transformed with corresponding plasmids. The cells were grown at 37 °C in LB medium with 100 μg of carbenicillin, or 50 μg of kanamycin, per ml. When the culture reached an optical density 600 nm (OD_600_) of 0.6-8, the culture was cooled to 22 °C, and 1 mM isopropyl-thiogalactopyranoside was added to induce the expression. The cells were harvested after additional growth for 12 hr at 22 °C, 180 rpm, and stored at −80 °C until use.

### Purification of recombinant proteins

Cell pellets were suspended in buffer (50 mM HEPES (pH 7.5), 5 mM MgCl_2_ containing lysozyme (hen egg white), DNase (TURBO, Thermo Fisher Scientific Inc.), and Complete^®^ protease inhibitor cocktail EDTA-free (Pierce)) that is twice the volume of cell pellet. The suspensions were stirred on ice for 2 hr and then sonicated 3x 30 sec on ice.

The cell lysates were centrifuged at 20,000 rpm, 4 °C, for 1 hr. The clear supernatant was incubated at 4 °C for 3 hr with Ni-NTA resin that has been equilibrated with buffer containing 10 mM imidazole. The resin slurry was then transferred to a glass Econo-column (Bio-Rad). The column was washed with the same buffer, and then performed a step-wise discontinuous imidazole gradient up to 500 mM imidazole. The fractions were analysed by SDS-PAGE. The protein fractions were pooled and concentrated using Vivaspin, and then purified by gel-filtration. The protein fractions were analysed by SDS-PAGE, pooled and concentrated using Vivaspin. The protein concentration was determined by BSA assay. The proteins were stored at −20 °C as 50% glycerol stock or −80 °C as 25% glycerol stock.

### Phylogenetic analysis

A seed multiple sequence alignment (MSA) of selected members of the HpcH_HpaI aldolase/citrate lyase family (PF03328) was built based on an alignment of the structures available in the PDB.[2] Structures were visualised and edited using Chimera and the multiple structural alignment was computed by Mustang.[3, 4] The PF03328 sequences in the full-alignment were downloaded from PFAM and aligned to the seed MSA using MAFFT E-INS-i --add option.[5] Partial and highly divergent sequences were removed and redundancy was reduced to 90% ID or less using Jalview.[6] The model of protein evolution used was LG+I+G+F with α of 1.64 and p-inv of 0.01, as selected by Prottest 3.4.[7] Bootstap repeats and consensus trees were generated with PHYLIP.[8] Pairwise distances were calculated with TREEPUZZLE *via* the puzzleboot script.[9] Distance trees were calculated with BIONJ.[10]

For the acyl-CoA lyase tree, sequences were aligned with MUSCLE and a maximum-likelihood tree was calculated with PhyML.[11, 12] Trees were drawn with Dendroscope 2.7.[13]

### Mycobacteria and culture conditions

*Mycobacterium tuberculosis* H37Rv (Mtb) was used in the studies, and all mycobacteria work was carried out in Containment 3 level facility. Mtb strains were grown at 37 °C in Middlebrook 7H9 medium (Sigma) supplemented with 10% albumin-dextrose-catalase supplement (ADC, Sigma) and 0.05% Tyloxapol (Sigma) or on Middlebrook 7H11 agar medium (Sigma) supplemented with 10% oleic acid-albumin-dextrose-catalase supplement (OADC, Sigma). Minimal medium is modified-7H9 or −7H10 medium without glycerol, L-glutamic acid, oleic acid, and dextrose. Chemically defined medium is modified-7H9 or −7H10 minimal medium with 10mM of one or more substrates (L-leucine, itaconate, glyoxylate, L-valine, L-isoleucine, succinate, α-ketoglutarate, citrate, acetate, oleate, and/or pyruvate).

### Mtb spotting culture

Mtb cultures were grown to OD_600_ of 1. A serial dilutions of 1:5. 2 μL of the OD_600_ of 1 and the diluted cultures were spotted on chemically defined-7H10 medium prepared with relevant test compounds. Plates were incubated at 37 °C. Growth phenotypes were observed, and images were taken on day: 25, 31, 53, and 195, post spotting.

### Mtb chemically defined medium growth culture

Mtb starter cultures were prepared by growing the cells to OD_600_ of 1. The starter cultures were used to inoculate 50 mL 7H9 medium supplemented with ADC and 0.05% Tyloxapol in roller bottles to OD_600_ of 0.05. On day 5, OD_600_ measurements were recorded and 10 mL of the cultures were transfer to 50-mL falcon tube. The cell pellets were collected after centrifugation at 2,000 rpm for 10 min at 4 °C, and washed with PBS. The washed cell pellets were suspended in conditioned-7H9 minimal medium with desired test compounds. The cultures were incubated at 37 °C on a rotator, with a fitted-rack to hold 50-mL falcon tubes, tilted at 60°, at 40 rpm. OD_600_ measurements were recorded on day: 3, 5, 7, 9, 12, 15, 19, and 26 post inoculation.

### Cell free protein extract preparation

Mtb cultures were grown at 37 °C in 7H9 medium supplemented with Tyloxapol and ADC to OD_600_ of 1. The cell pellets were collected after centrifugation at 2,000 rpm for 10 min at 4 °C, washed with HEPES (50 mM, pH 7.5, 5 mM MgCl_2_). The washed pellets were suspended in HEPES buffer and transferred to O-ring screw cap tube with glass beads (150-212 μm, acid washed, Sigma), and ribolysed for 2 × 30 sec with cooling between the two cycles. The clear supernatants were collected after centrifugation at 13,000 rpm for 10 min at 4 °C, and filtered twice through 0.2 μm membrane. The total protein concentration was determined by BCA assay (Pierce).

### Syntheses of CoA-thioesters

Citramalyl-CoA, malyl-CoA, and β-methylmalyl-CoA were synthesised enzymatically with MMclyase, as described by Zarzycki *et al.*[14] Citryl-CoA was synthesised enzymatically with inactivated citrate lyase from *Klebsiella pneumoniae* (Sigma), as described by Buckel *et al.*[15] The CoA-thioesters were analysed by HPLC, and their identities were confirmed by high resolution mass spectrometry.

### Murine aerosol *M. tuberculosis* infections

C57BL/6J (WT) mice were bred and maintained under specific pathogen-free conditions at The Francis Crick Institute, Mill Hill Lab. Animal studies and breeding were approved by the Francis Crick Institute ethical committee and performed under U.K Home Office project license PPL 70/8045. Infections were performed in the category 3 animal facility. For mouse infections *M. tuberculosis* strains were cultured in Middlebrook 7H9 broth containing ADC to an OD_600_ of 0.6. From this, an infection sample was prepared to enable delivery of 100 colony forming units (CFUs)/mouse lung using a modified Glass-Col aerosol infection system. Infection was monitored by assessing homogenised lungs from infected mice at defined periods post-infection. Bacterial CFUs were determined by plating serial dilutions of homogenates on duplicate Middlebrook 7H11 containing OADC. Colonies were counted 2-3 weeks after incubation at 37 °C. The data at each time point are the means of 5 mice/group +/− SEM.

### Gene synthesis

Gene sequences (*PA0883*, *PA2011*, and *MMClyase*) were codon-optimized for *Escherichia coli* expression, synthesised, and then cloned (NdeI-BamHI) into pET-16b by GenScript (Piscataway, NJ).

### Other techniques

DNA sequencing was performed by GATC Biotech (Konstanz, Germany). Protein concentration was determined by bicinchoninic acid (BCA) assay (Pierce BCA Protein Assay Kit), using bovine serum albumin as the standard.

### Data analysis

Data were processed by GraphPad PRISM 7.02, Agilent Qualitative Analysis B.07.00, MestReNova 12.0.1-20560, PyMOL.

### LC/MS metabolomics

Methods are as described by Larrouy-Maumus *et al.*[16]

### HPLC-based activity assays

Rv2498c or cell free protein extract (CFPE) reactions contained 50 mM HEPES (pH 7.5), 5 mM MgCl_2_, and specified concentration of substrate (usually in the 10-50 μM range). Reactions were initiated by the addition of enzyme or CFPE, and incubated at 37 °C for a given amount of time. The reactions were terminated on ice and quenched with 1 μL of 1M HCl per 10 μL reaction volume (or without quenching, and directly used for HPLC analysis), centrifuged at 13,000 rpm for 10 min at 4 °C, and the clear supernatants were used for HPLC analysis. HPLC-based methods were used to directly detect the CoAs with UV absorption at 260 nm. An Agilent 1260 Infinity (Santa Clara, CA) HPLC apparatus was used, equipped with G1311B 1260 QuatPump, G1316A 1260 TCC, G1264C 1260 FC-AS, G1267E 1260 HiP ALS, and G4212B 1260 DAD.

Each CoA-thioester was detected as a single peak after being separated by HPLC with a Poroshell 120 EC-C18, 2.7 μm, 4.6 × 50 mm column (Agilent) using the following elution condition: 1-min isocratic elution at 2% acetonitrile in buffer, followed by a 10-min linear gradient of 2-20% acetonitrile, with 2-min isocratic elution at 20% acetonitrile in buffer, and then back to 1-min isocratic elution at 2% acetonitrile in buffer at a flow rate of 0.5 ml/min.

The buffer used was either 40 mM ammonium formate, pH 6.8 or 40 mM potassium phosphate, pH 7.0. For more hydrophobic CoA-thioesters, 95% instead of 20% acetonitrile was used for the gradient. When necessary, CoA-thioesters were quantified by a calibration curve generated from the authentic Acetyl-CoA standard (Sigma-Aldrich) stock solutions (5, 15, 25, 50, 100, 150, 250 μM).

### ^1^H Proton nuclear magnetic resonance spectroscopy

Samples were prepared in D_2_O or D_2_O phosphate buffer (0.1 M, pD = 7.2). Proton nuclear magnetic resonance (δH) spectra were recorded on Bruker Avance III HD 400 (400 MHz), Bruker Avance III 600 (600 MHz), or Bruker Avance III HD 800 (800 MHz). All chemical shifts were quoted on δ-scale in ppm, with residual solvent as internal standard.

### Construction of Mtb *rv2498c*KO mutant and complements

The construction of Rv2498c knockout followed methods described by Parish et al.[17] Briefly, an unmarked in frame deletion of the *Rv2498c* gene was made by amplifying 1.5 kb of flanking sequence from Mtb H37Rv with Phusion taq (upstream of Rv2498c: 5’-GCAGATCTTCGGCGGCCATCGCGTCGTA-3’, 5’-GCTCTAGAACCTCCGAATGAGGGCGCAG-3’; downstream of Rv2498c: 3’ for: 5’-GCTCTAGAACGCAGGTTCATTGCGCCTC-3’, 5’-GCAGATCTTGGTACTTGAGGAGCTGGGC-3’), and cloned into PCR4blunt (Invitrogen). The inserts digested with *BglII* and *XbaI*, and the fragments were ligated together using T4 DNA ligase. The ligation product was amplified (5’-GCAGATCTTCGGCGGCCATCGCGTCGTA-3’, 5’-GCTCTAGAACGCAGGTTCATTGCGCCTC-3’), digested with *BglII* and cloned into the *BamH1* site of p2NIL. The *PacI* fragment of pGOAL17 containing the *lacZ* and *sacB* genes was cloned into the *PacI* site of the resulting plasmid to make the final *rv2498c*KO construct plasmid. Competent Mtb H37Rv was electroporated with the knockout construct plasmid, and single crossovers were selected on 7H11 plates containing kanamycin and X-gal. The blue colonies were then streaked on 7H11 plates and then double crossovers were selected on 7H11 plates containing sucrose and X-gal, the resulting white colonies were screened for double crossovers. Southern blot was performed to confirm a knockout of Rv2498c.

For the complement, *rv2498c* gene fragment was amplified from Mtb H37Rv genomic DNA by PCR (5’-AACAGAAAGGAGGTTAATAATGAACCTGCGTGCCGCC-3’, 5’-TTAGCTAAAGCTTATTTAAATTCATTCGGAGGTGGCTTCCCCG-3’) and assembled with pML1335 integrative vector PCR fragment (5’-ATTTAAATAAGCTTT AGCTAATTAATT GGGGACCCTAGAGGTC-3’, 5’-TATTAACCTCCTTTCTGTTAATTAAGCATGCGGATCGT-3’) by Gibson assembly (NEB) to create pML1335-P_smyc_Rv2498c. A second Rv2498c complement plasmid was created with *Pimyc* constitutive promoter. Similarly as before, *Pimyc* gene sequence was amplified from pMC2m by PCR (5’-TGTTTAAACTCTAGAAATATTGGATCGTCGCACCGGGTTAAG-3’, 5’-CTTTCTGTTAATTAAGCATGCGGATCCGTGGCAGGAGGGTC-3’) and assembled with pML1335-P_smyc_Rv2498c PCR fragment without *Psmyc* sequence (5’-GCATGCTTAATTAACAGAAAGGAGGTTAATA-3’, 5’-AATATTTCTAGAGTTTAAACACTAGTCGAGAAAAAAAAAAGCGC-3’) by Gibson assembly (NEB) to create pML1335-P_imyc_Rv2498c. Competent Mtb:*rv2498c*KO were electroporated with pBS-Int (for integrase) with pML1335-P_smyc_Rv2498c and pML1335-P_imyc_Rv2498c, and then transferred to 5 mL 7H9 medium with supplement (ADC), and Tyloxapol, and left standing overnight at 37 °C. The electroporated cells were plated on 7H11 medium with supplement (OADC) containing 100 mg/mL Hygromycin B to select for complemented colonies. Complementation were checked by PCR (same primers used in gene amplification noted above) with genomic DNA obtained by InstaGene Matrix (Bio-Rad).

### Metabolite preparation

Mtb strains were grown at 37 °C in 7H9 medium supplemented with Tyloxapol and ADC to OD_600_ of 1. Drop by drop, total of 1-mL of the culture was transferred to a membrane disc under vacuum to collect the cells on the membrane. The membrane disc containing the cells was transferred to 7H10 medium supplemented with OADC, and incubated for 5-6 days until sufficient cell mass accumulation. The membrane disc containing abundant cell mass was transferred to conditioned-7H10 minimum medium supplemented relevant test substrate, and incubated for an additional 17 hr maximum to avoid doubling of the cells. After the cells were incubated with the test substrate, the cells were scrapped and transferred to an O-ring screw cap tube with glass beads (150-212 μm, acid washed, Sigma), with the addition of 0.5-0.7 mL cold acetonitrile:methanol:water (2:2:1), and ribolysed for 2 × 30 sec with cooling between the two cycles. The clear supernatant was collected after centrifugation at 13,000 rpm for 10 min at 4 °C, and filtered twice through 0.2 μm membrane. The filtered supernatant was analysed by LC/MS, or stored at −80 °C until further use.

### Crystallization and Data Collection

Preparations of Rv2498c containing various ligands crystallized under two general conditions, acetate pH 7 and sulphate (or phosphate) pH ~5, yielding different crystal forms (R32 or C2, respectively). Crystals were grown using the sitting drop vapour diffusion method at room temperature. The initial protein solution contained Rv2498c at a concentration of 15 mg/mL in 50 mM HEPES pH 7.4 and 300 mM NaCl. For final crystallization conditions for the six complexes, see **Supplementary Information**.

Prior to data collection, all crystals were transferred to cryoprotectant solutions composed of their mother liquors supplemented with 20% glycerol and flash-cooled in a nitrogen stream at 100K. X-ray diffraction data were collected by LRL-CAT staff at APS beamline 31-ID-D (Advanced Photon Source, Argonne National Laboratory, Argonne, IL) on a Rayonix 225-HE detector (Rayonix). Diffraction intensities were integrated, scaled and merged with the programs DENZO and SCALEPACK.[18] Data collection statistics are given in **Supplementary Table 2**.

### Crystallographic Structure Determination and Refinement

See **Supplementary Information** for the complete structure determination and refinement of other ligands.

*Rv2498c‧Mg^2+^‧CoA‧Acetoacetate:* The complex structure of Rv2498c with Mg^2+^, acetoacetate and CoA was determined by molecular replacement with PHENIX using the complex of Rv2498c with Mg^2+^ and acetoacetate, determined earlier, as a search model.[19, 20] Several rounds of automated and manual building and refinement, with COOT, ARP/wARP, and PHENIX, respectively, converged with R_work_=0.217 and R_free_=0.277 at 2.04Å resolution.[20–22] The final model consists of Rv2498c residues 3-269 in all three chains of the trimeric asymmetric unit (the N-terminal cloning artefacts, affinity tag, and residues 1, 2, and 270-273 were not observed). A magnesium ion, coordinated acetoacetate, and CoA molecule were well ordered in the active sites of three of the Rv2498c subunits and were included in the refined model. Additionally, two glycerol molecules and three sulphate ions were located and included in the final model. The coordinates and structure factors have been deposited in the PDB as entry 6AS5.

*Rv2498c‧Mg^2+^‧Citramalyl-CoA‧Pyruvate:* The complex structure of Rv2498c with Mg^2+^, pyruvate, and citramalyl-CoA was determined by molecular replacement with PHENIX using the complex of Rv2498c with Mg^2+^ and pyruvate, determined earlier, as a search model.[19, 20] Several rounds of manual building and refinement, with COOT and PHENIX, respectively, converged with R_work_=0.191 and R_free_=0.233 at 1.83Å resolution.[20–22] The final model consists of Rv2498c residues in three chains of the trimeric asymmetric unit (chain A, 2-268; chain B, 1-268; chain C, 1-267; the N-terminal cloning artefacts, affinity tag, and residues 1 in chain A, 269-273 in chains A and B, and 268-273 in chain C were not observed). A magnesium ion was located in each active site; two citramalyl-CoA molecules (presumably arising as the product of enzyme mediated reaction between pyruvate and acetyl-CoA both of which were present in the crystallization milieu) were well ordered in the active sites of chains A and B and were included in the model. The magnesium ion in the active site of chain C was coordinated by a pyruvate molecule. Additionally, two glycerol molecules and three phosphate ions and five chlorides were located and included in the final model. The coordinates and structure factors have been deposited in the PDB as entry PDB entry 6AQ4.

## SUPPLEMENTARY INFORMATION

**Figure S1.**
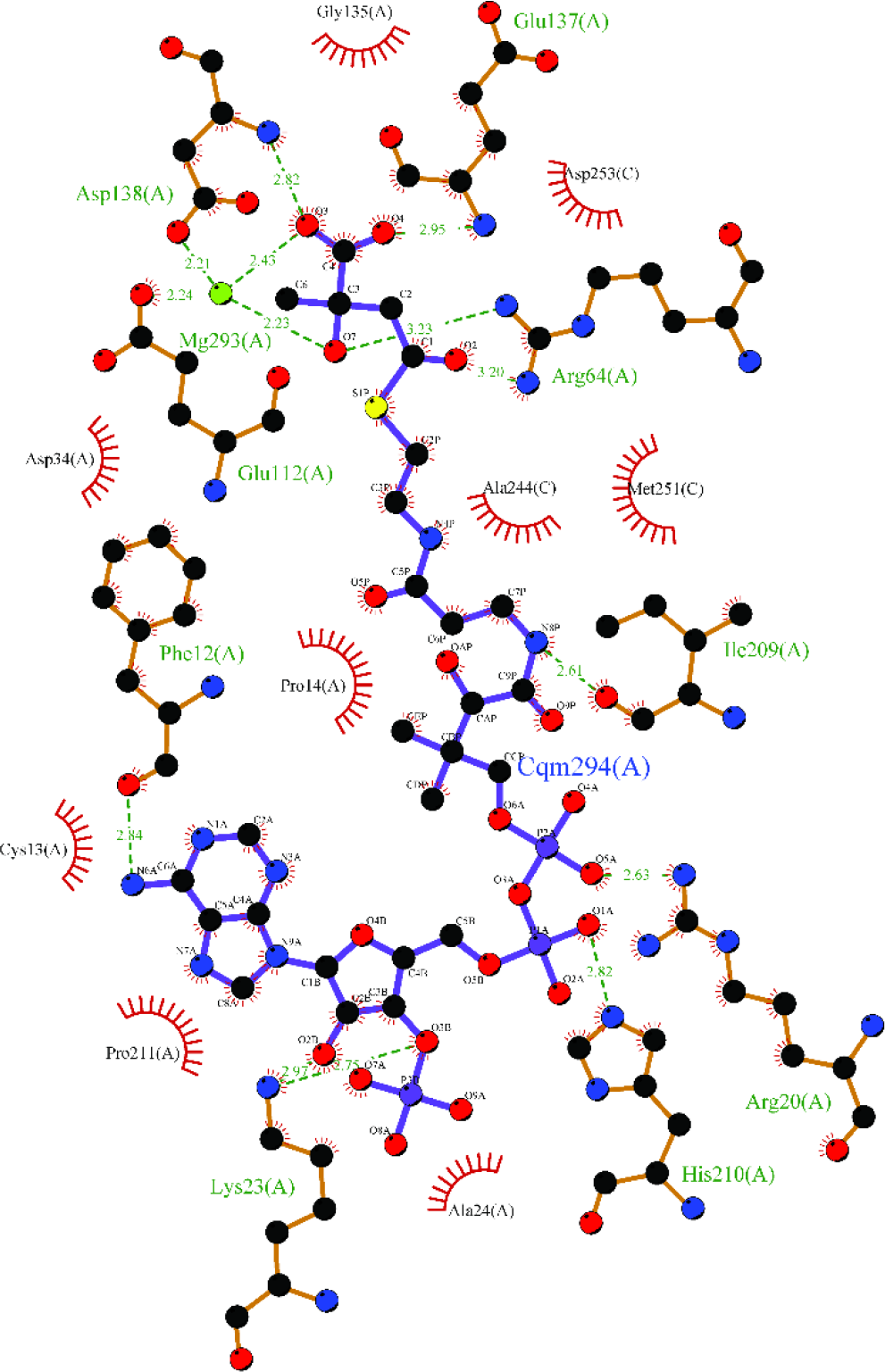
Stick representation of Rv2498c-ligand complex. Rv2498c-ligand complex of active site residues responsible for interaction with magnesium metal and (*S*)-citramalyl-CoA. Image was generated with Ligplot+.

**Figure S2.**
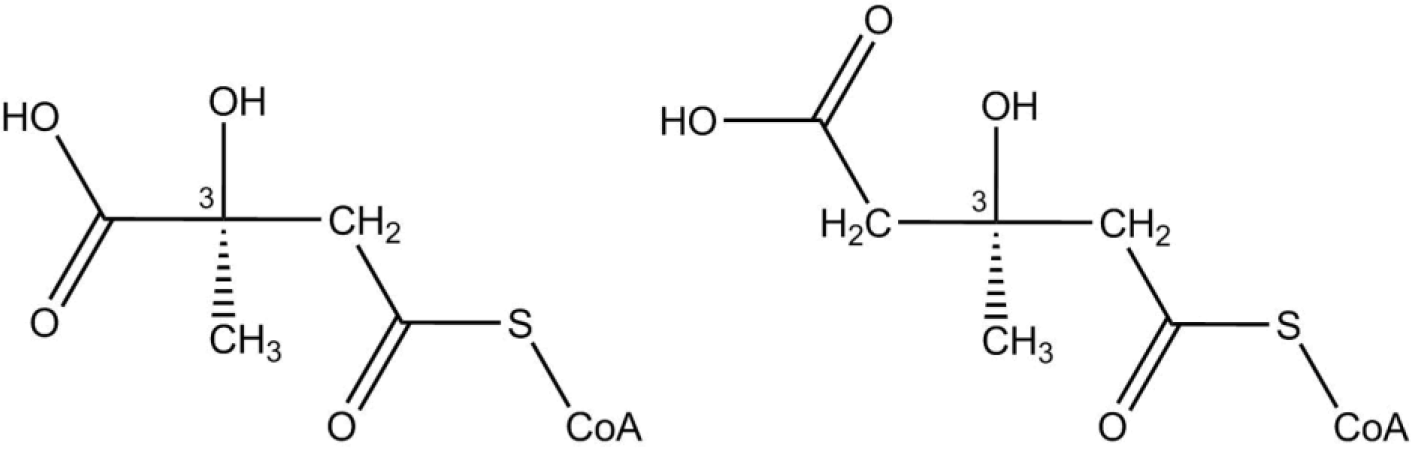
Stereochemistry of 3-carbon position of β-hydroxyl-acyl-CoA thioesters. Stereochemistry at the 3 carbon positions of *S*-citramalyl-CoA (left) and *R*-HMG-CoA (right).

**Figure S3.**
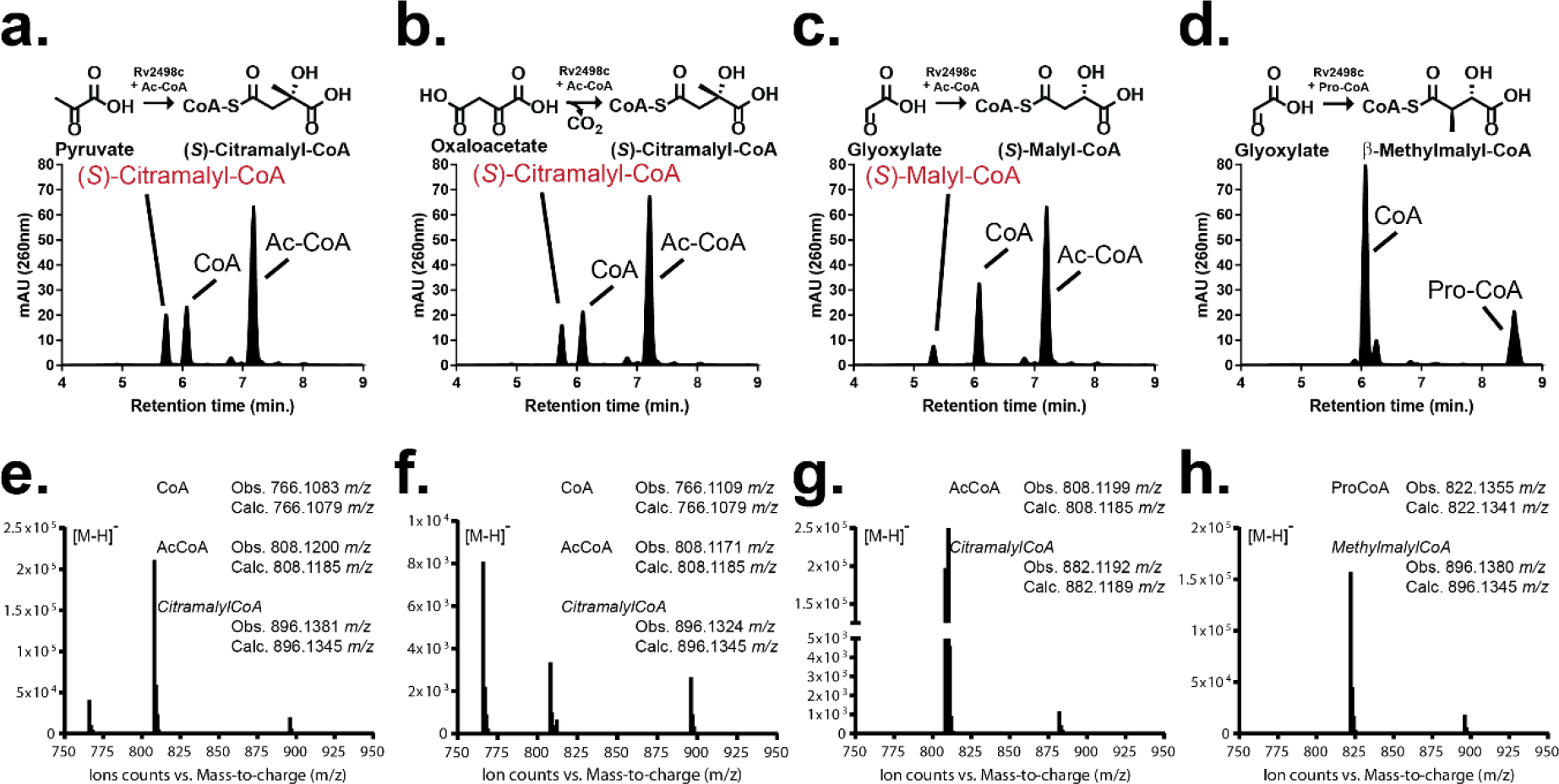
Rv2498c synthase acyl-CoA thioester reactions. HPLC chromatograms of Rv2498c reverse reactions with keto-acids and acyl-CoAs after 60 min at 37 °C. A reaction mixture typically contains 150 μM of the acyl-CoA and the 30 mM of keto-acid (molar ratio of 1:200, respectively), and 0.5 μM Rv2498c. (**a**) Rv2498c synthesises citramalyl-CoA from pyruvate and acetyl-CoA (**b**) Rv2498c synthesises citramalyl-CoA, instead of citryl-CoA, from oxaloacetate and acetyl-CoA, due to spontaneous decarboxylation of oxaloacetate to pyruvate. (**c**) Rv2498c synthesises malyl-CoA from glyoxylate and acetyl-CoA. (**d**) Rv2498c synthesises methylmalyl-CoA from glyoxylate and propionyl-CoA. (**e-h**) MS spectra of Rv2498c synthase reaction mixtures. A concentrated substrates (acyl-CoA:keto-acid, 0.3 mM: 60 mM) to enzyme (0.5 μM) ratio reaction mixture was prepared to observe reaction acyl-CoAs.

**Figure S4.**
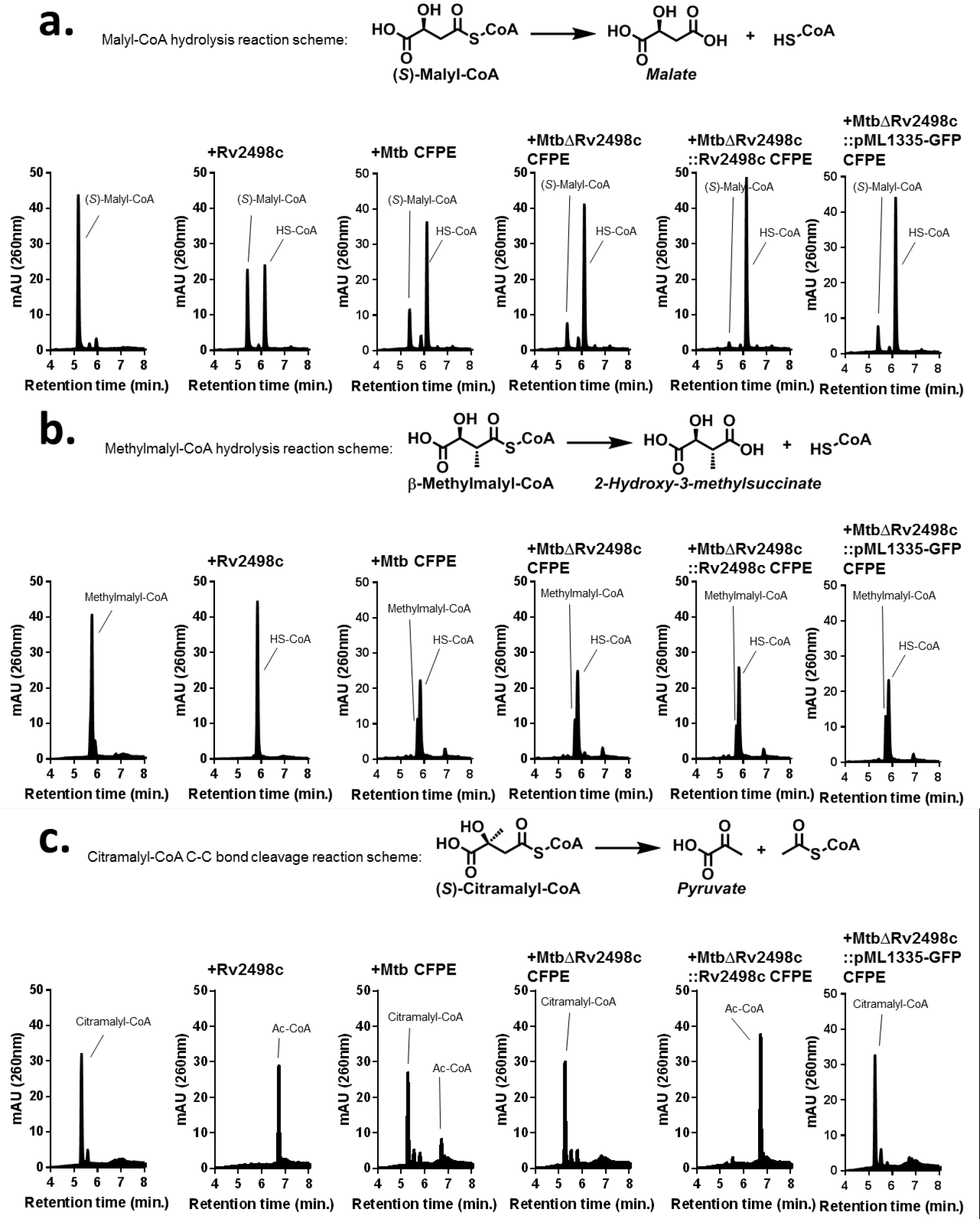
Rv2498c hydrolysis of Malyl-CoA and Methylmalyl-CoA, and C-C bond cleavage of Citramalyl-CoA. HPLC chromatograms of in vitro Rv2498c reactions. (**a**) Rv2498c hydrolysis (*S*)-malyl-CoA to HS-CoA and malate. (**b**) Rv2498c hydrolysis β-methylmalyl-CoA to HS-CoA and 2-hydroxy-3-methylsuccinate (methylmalate). (**c**) Rv2498c C-C bond cleavage of (*S*)-citramalyl-CoA to acetyl-CoA and pyruvate.

**Figure S5.**
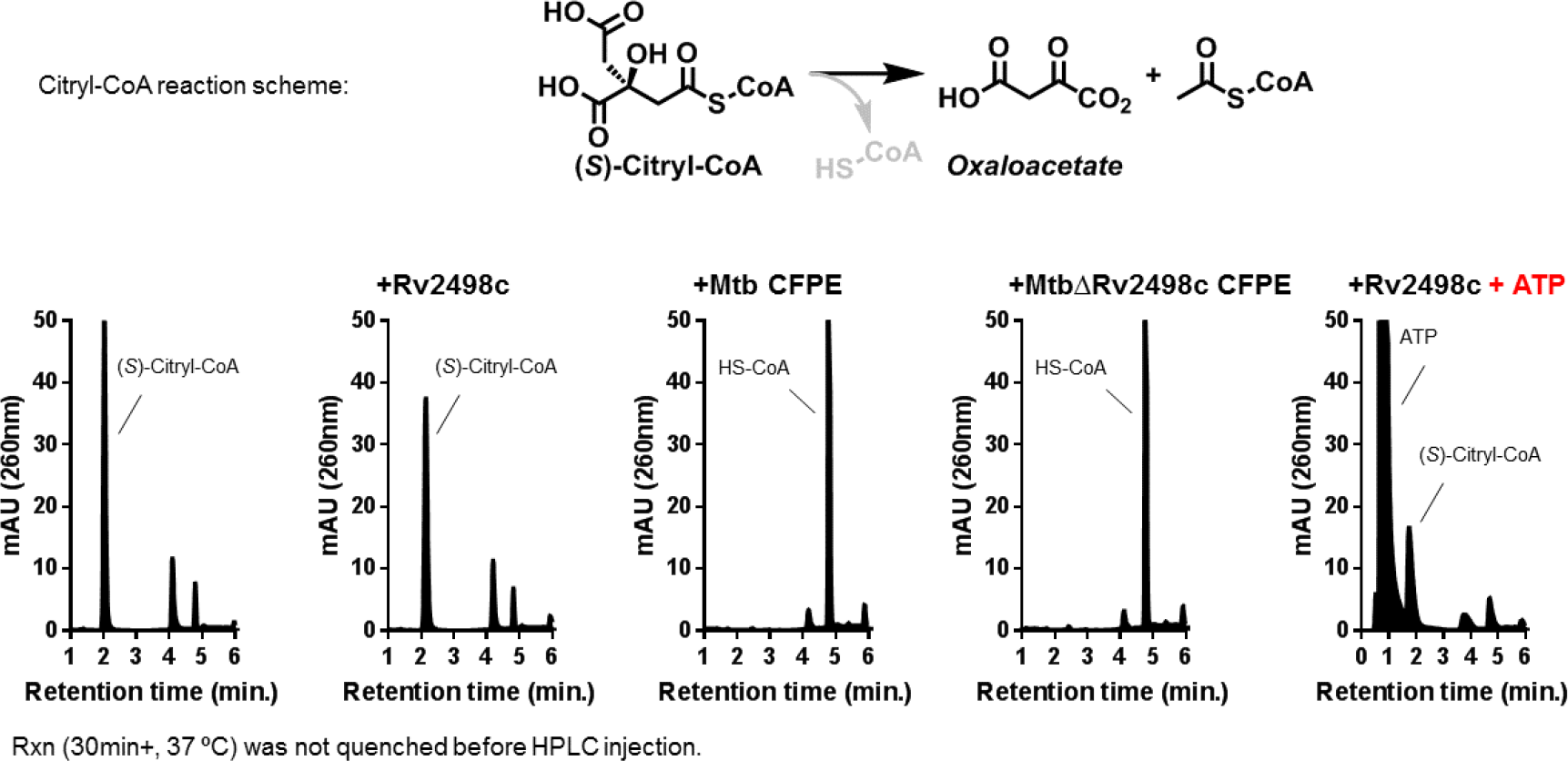
Investigation of Citryl-CoA as a substrate for Rv2498c. (*S*)-Citryl-CoA, synthesised from inactivated citrate lyase, was used as the substrate to confirm Rv2498c does not consume (*S*)-citryl-CoA. Even with the addition of ATP with (*S*)-citryl-CoA, Rv2498c does not consume (*S*)-citryl-CoA. Rv2498c is neither an ATP-independent nor an ATP-dependent citryl-CoA lyase. (*S*)-citryl-CoA is hydrolysed by both WT and *rv2498c*KO CFPEs to suggest the hydrolysis is carried out by another enzyme, such as a citrate synthase.

**Figure S6.**
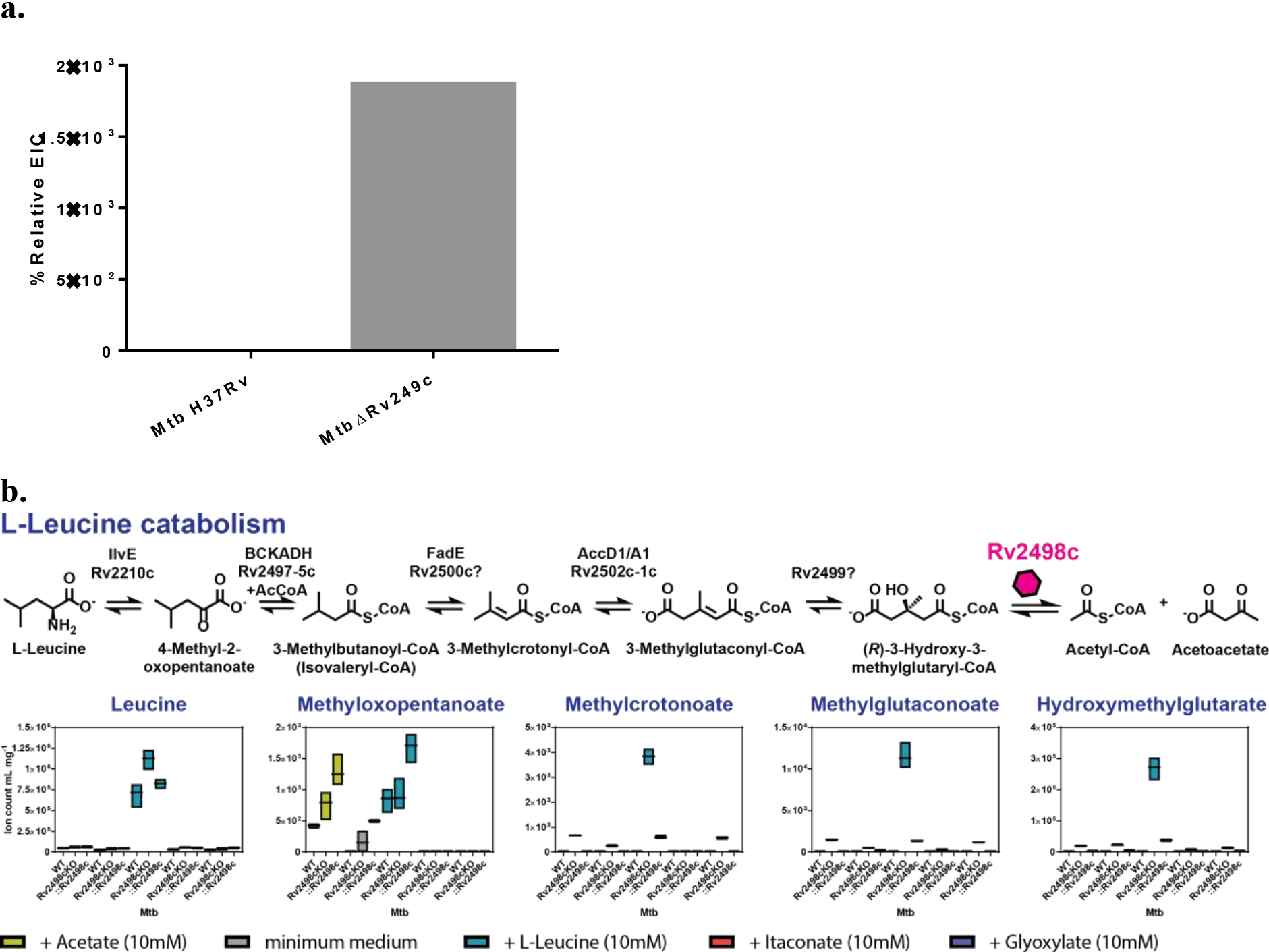
Metabolic profile from Mtb growth in L-leucine chemically defined medium. (**a**) The cell culture medium from L-leucine growth from Mtb WT and *rv2498c*KO by LC-MS, and found the *rv2498cKO* strain did indeed accumulate HMG, but not WT. Metabolite profile shows quantitative measurements of metabolite in growth medium with added substrate (10 mM). (**b**) The accumulation of hydroxymethylglutarate (HMG) and other L-leucine catabolic pathway keto-acid intermediates in the *Rv2498cKO* was similarly detected in extracted metabolite samples by LC-MS. Metabolic profiles show quantitative measurements of metabolite from filter culture growth on chemically defined media with added substrate (10 mM). The data are shown as mean values ± SD from three biological replicates.

**Figure S7.**
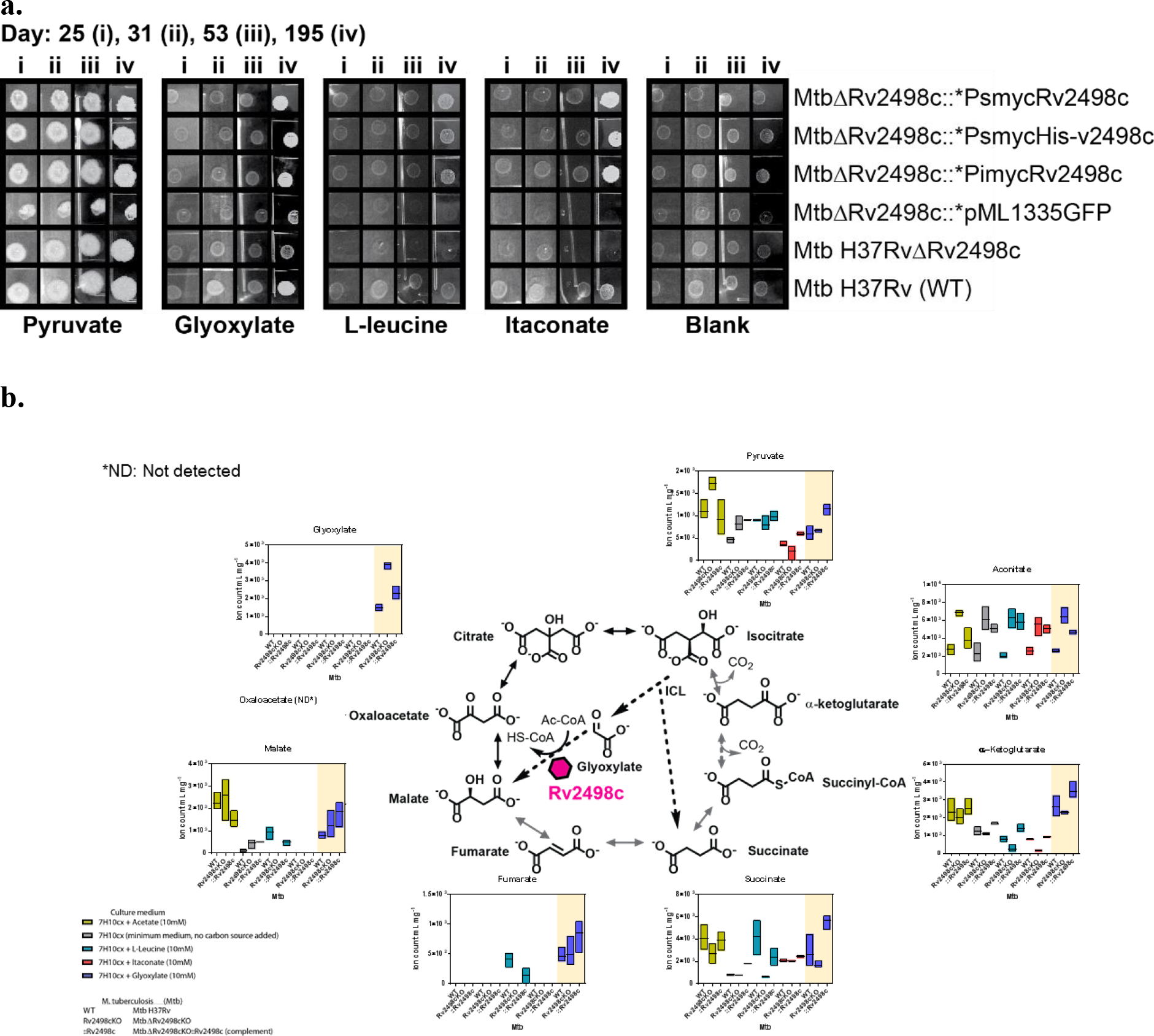
Mtb spotting experiments, and TCA metabolic profiles from Mtb growth in glyoxylate chemically defined medium. (**a**) Images from spotting experiment from different Mtb strains in chemically defined medium. The results are representative of three independent experiments. (**b**) Metabolic profile from extracted metabolite samples of 17 hr Mtb filter growth on chemically defined medium (glyoxylate, acetate, L-leucine, and itaconate). Metabolic profiles show quantitative measurements of metabolite from filter culture growth on chemically defined media with added substrate (10 mM). The data are shown as mean values ± SD from three biological replicates.

**Figure S8.**
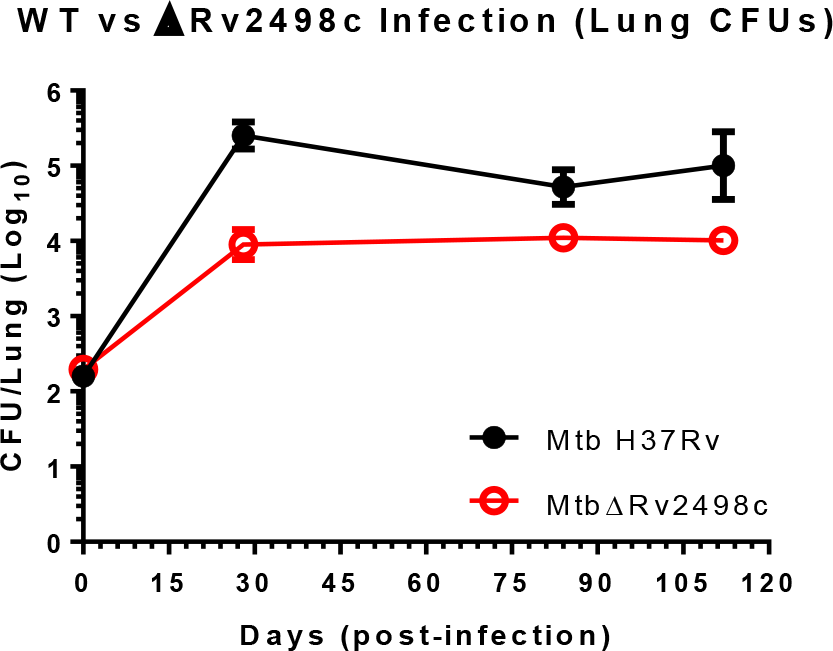
Mice lung infection study with Mtb H37Rv and Rv2498c mutant. Rv2498cKO displays 1 log10 reduction in colony forming units (CFUs) in lung at day 28, 84, 112 post infection, compared to parent strain Mtb H37Rv.

**Figure S9.**
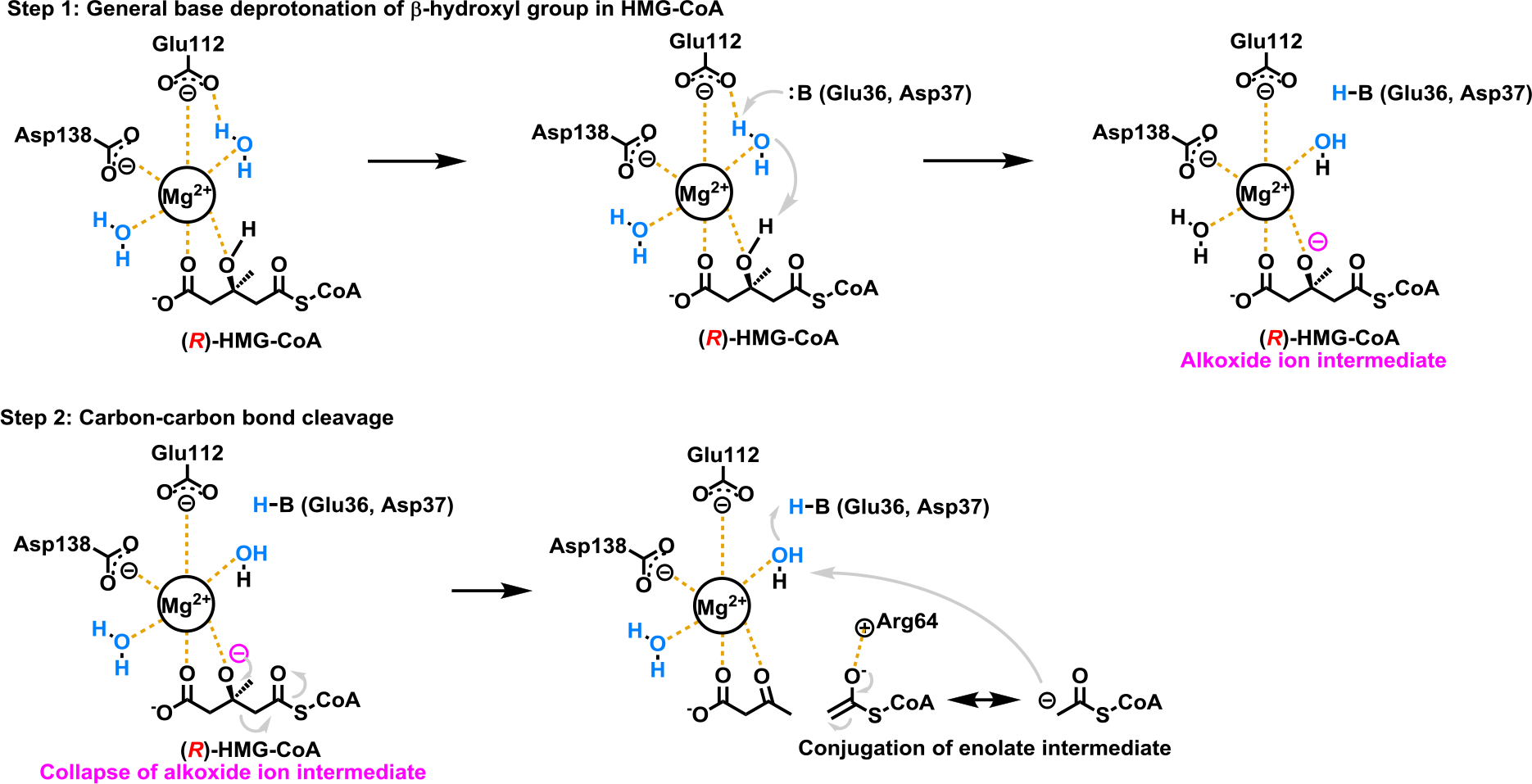
A proposed mechanism for Rv2498c C-C cleavage of β-hydroxyl-acyl-CoA thioesters. The first step in the reaction is the base catalysed deprotonation of the β-hydroxyl group of HMG-CoA *via* metal-bound water-mediated proton abstraction. Based on our structures, the general base is either Glu36 or Asp37, both at close proximity (less than 3 angstroms) to the water molecules. This first step generates an alkoxide intermediate. The second step is the C-C bond cleavage as a result from the collapse of the alkoxide resulting in the two reaction products: acetoacetate and acetyl-CoA. The Arg64 is suggested to partially involve in the enolisation of the acetyl-CoA intermediate. The general base is likely regenerated by the metal-bound water-mediated proton abstraction.

**Table S1.**
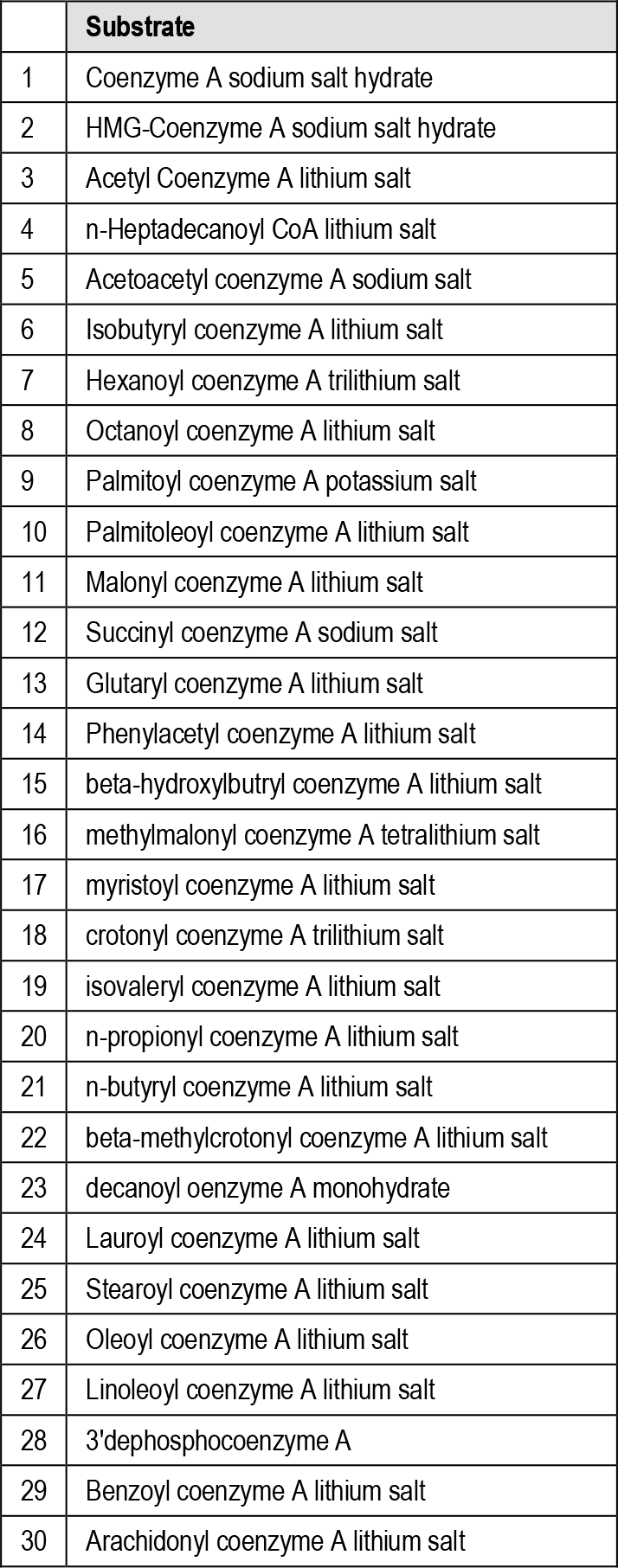
CoA-thioesters.

**Table S2.**
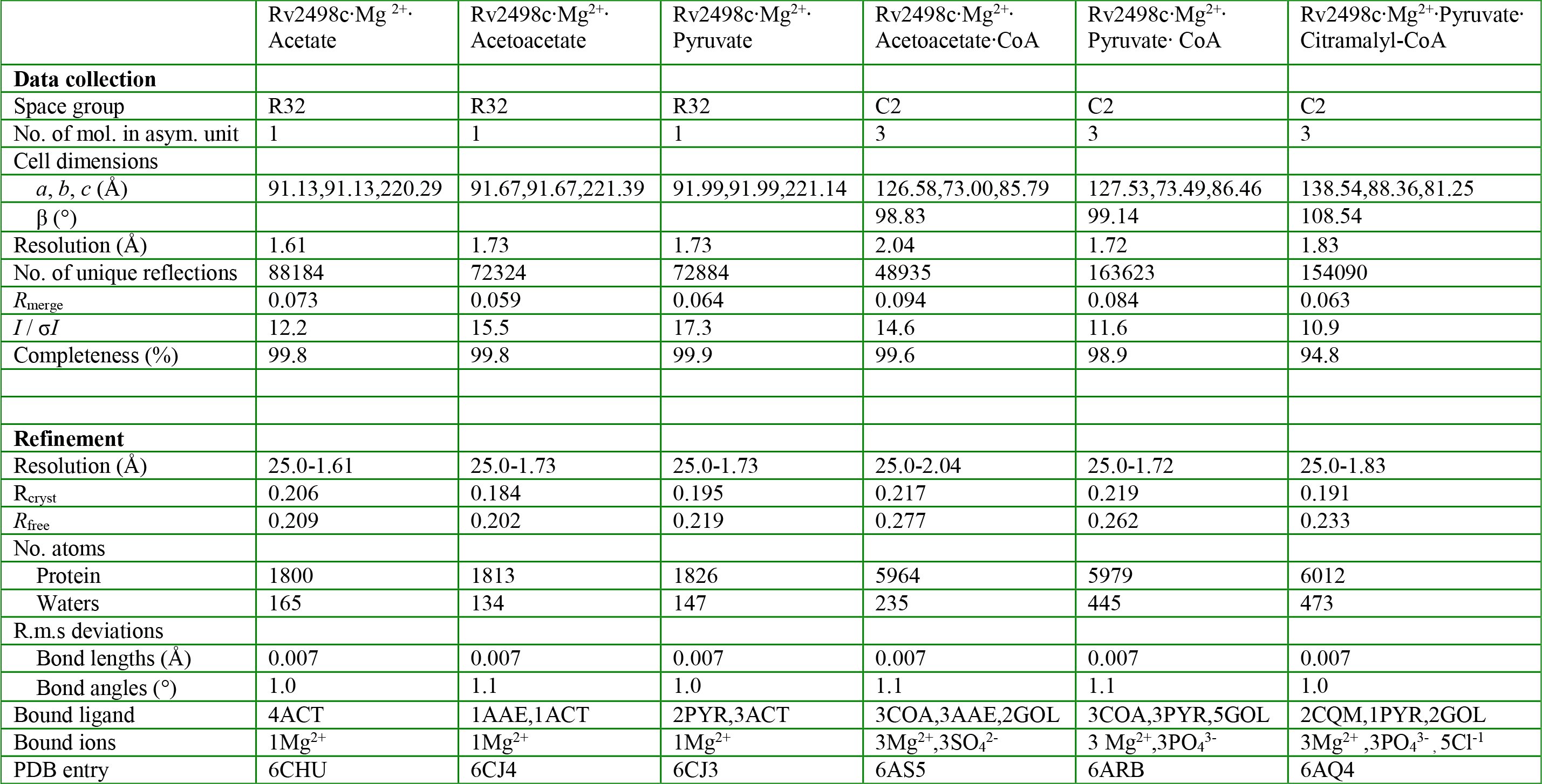
X-ray data collection and refinement statistics for complexes of Rv2498c from *Mycobacterium tuberculosis*.

### Crystallization Conditions

For final crystallization conditions for the six complexes

1. Rv2498c‧Mg^2+^‧CoA‧Acetoacetate: to a protein solution containing 10 mM MgCl_2_, 40 mM pyruvate, and 40 mM Acetyl-CoA was added precipitant contained 2 M ammonium sulfate pH 5.5. Crystals appeared in one week and exhibited diffraction consistent with the space group C2 with three molecules of the complex per asymmetric unit.
2. Rv2498c‧Mg^2+^‧Pyruvate‧Citramalyl-CoA: to a protein solution containing 20 mM MgCl_2_, 100 mM pyruvate, and 20 mM acetyl-CoA was added a precipitant containing 0.4 M ammonium phosphate pH 4.2. For this sample crystals appeared in three weeks and exhibited diffraction consistent with the space group C2 with tree molecules of the complex per asymmetric unit.
3. Rv2498c‧Mg^2+^‧Acetate: to a protein solution containing 10 mM MgCL_2_ was added a precipitant containing 1.0 M sodium acetate pH 7.0. Crystals appeared in 4-5 days and exhibited diffraction consistent with the space group R32, with one molecule of the complex per asymmetric unit.
4. Rv2498c‧Mg^2+^‧Acetoacetate: to a protein solution containing 10 mM MgCl_2_ and 2 M acetoacetate was added a precipitant containing 1.0 M sodium acetate pH 7.0. Crystals appeared in 3 days and exhibited a diffraction pattern consistent with space group R32, with 1 molecule of the complex per asymmetric unit.
5. Rv2498c‧Mg^2+^‧Pyruvate: to a protein solution containing 10 mM MgC1_2_ and 2 M pyruvate was added a precipitant contained 1.0 M sodium acetate pH 7.0. Crystals appeared in 2 days and exhibited a diffraction pattern consistent with space group R32 with one molecule of the complex per asymmetric unit.
6. Rv2498c‧Mg^2+^‧CoA‧Acetoacetate: to a protein solution containing 10 mM MgCl_2_, 40 mM acetoacetate, and 40 mM Acetyl-CoA was added a precipitant contained 2 M ammonium sulfate pH 5.5. Crystals appeared in one week and exhibited diffraction consistent with the space group C2 with 3 molecules of the complex per asymmetric unit.

### Crystallographic Structure Determination and Refinement

*Non-CoA ligands:* The structures of three of the complexes, Rv2498c‧Mg^2+^‧Acetate, Rv2498c‧Mg^2+^‧Acetoacetate, and Rv2498c‧Mg^2+^‧Pyruvate, were solved by molecular replacement using PHENIX with the apo structure of Rv2498c from *M. tuberculosis* (PDB ID 1U5H; residues 1-221) as a search model. [1, 2]

Automatic model building with ARP/wARP was followed by iterative cycles of manual model building and refinement, performed with COOT and PHENIX, respectively.[2–4] Refinement of the complexes converged at the following values: Rv2498c‧Mg^2+^‧Acetate: R_work_=0.206, R_free_=0.209 at 1.61Å resolution (PDB entry 6CHU); Rv2498c‧Mg^2+^‧Acetoacetate: R_work_=0.184, R_free_=0.202 at 1.73Å resolution (PDB entry 6CJ4); Rv2498‧Mg^2+^‧Pyruvate: R_work_=0.195, R_free_=0.219 at 1.73Å resolution (PDB entry 6CJ3). Refinement statistics are given in Table S2.

The final models of 6CHU, 6CJ3 and 6CJ4 consist of the following:

1) 6CHU – amino acids 1-221 and 250-265 of Rv2498c and one His of the N-terminal affinity tag. All other cloning artifacts along with amino acids 222-249 and 266 to 273 were not observed in the electron density, presumably due to disorder. One magnesium ion and four acetate molecules were also well ordered in the structure and were included in the final model.
1) 6CJ4 – amino acids 1-224 and 251-265 of Rv2498c. All cloning artifacts along with amino acids 225-250 and 266 to 273 were not observed in the electron density, presumably due to disorder. One magnesium ion, one acetate, and an acetoacetate molecule were also well ordered in the structure and were included in the final model.
1) 6CJ3 – amino acids 2-224 and 250-265 of Rv2498c. All cloning artifacts along with amino acids 1, 225-249 and 266 to 273 were not observed in the electron density, presumably due to disorder. One magnesium ion, three acetates, and two pyruvate molecules were also well ordered in the structure and were included in the final model.

*Rv2498c‧Mg^2+^‧CoA‧Pyruvate:* The complex structure of Rv2498c with Mg^2+^, pyruvate and CoA was solved by molecular replacement with PHENIX using the complex of Rv2498c with Mg^2+^ and pyruvate, determined earlier, as a search model.[1, 2] Several rounds of building and refinement, with COOT and PHENIX, respectively, converged with R_work_=0.219 and R_free_=0.262 at 1.72Å resolution.[2–4] The final model consists of Rv2498c residues 2-269 in each of three chains of the trimeric asymmetric unit (the N-terminal cloning artifacts, affinity tag, and residues 1 and 270-273 were not observed). A magnesium ion, coordinated pyruvate, and CoA molecule were observed in the active sites of three of the Rv2498c subunits and were included in the refined model. Additionally, five glycerol molecules and three phosphate ions were located and included in the final model. The coordinates and structure factors have been deposited in the PDB as entry 6ARB.

Final crystallographic details for all Rv2498c X-ray structures are provided in **Supplementary Table 2**.

